# Low-barrier hydrogen-bond powers long-range radical transfer in the metal-free ribonucleotide reductase

**DOI:** 10.64898/2026.03.06.710006

**Authors:** Abhishek Sirohiwal, Juliane John, Yury Kutin, Rohit Kumar, Federico Baserga, Vivek Srinivas, Hugo Lebrette, Maximilian C. Pöverlein, Ana P. Gamiz-Hernandez, Joachim Heberle, Müge Kasanmascheff, Martin Högbom, Ville R. I. Kaila

## Abstract

Ribonucleotide reductases (RNRs) catalyze the conversion of ribonucleotide (RNA) to deoxyribonucleotide (DNA) building blocks initiated by a long-range (>30 Å) proton-coupled electron transfer (PCET) by mechanistic principles that remain much debated. By combing multiscale quantum and classical simulations with directed mutagenesis, x-ray crystallography, and vibrational and electron paramagnetic resonance spectroscopy, we elucidate here the molecular principles underlying how metal-free RNRs initiate the long-range PCET process by creating a highly stable DOPA initiator radical. We show that DOPA• is redox-tuned by a low-barrier hydrogen bond (LBHB), with a delocalized proton that provides the catalytic power for the ribonucleotide reduction. We find that the LBHB couples to an extended hydrogen-bonded network, with distant mutations resulting in the loss of radical formation, and providing key molecular insight into the long-range radical transport mechanism in RNRs. On a general level, our findings support the direct involvement of LBHB in protein chemistry and the importance of quantum effects in enzyme catalysis.

**Significance Statement:** Ribonucleotide reductases (RNRs) are ancient enzymes responsible for the synthesis of deoxyribonucleotides from ribonucleotides. RNRs catalyze this reaction *via* a long-range proton-coupled electron transfer (PCET) process, involving the formation of a stable protein radical. Yet, despite decades of detailed structural, biochemical, spectroscopic and computational studies, the mechanistic principles of this process remain unclear and much debated. Here, we show that metal-free RNRs power the reduction of RNA building blocks by a highly stable organic DOPA initiator radical, arising from a unique low-barrier hydrogen bonding (LBHB) network that enables the radical transport by strong redox-tuning effects. Our findings reveal mechanistic principles underlying the elusive PCET reactions of metal-free RNRs, and provide evidence for the involvement of quantum effects in enzyme catalysis.

## Introduction

Proton-coupled electron transfer (PCET) reactions are central charge transfer processes in biology (1, 2). In cellular respiration, membrane-bound enzymes rely on PCET reactions to oxidize nutrients and create proton currents that power the synthesis of adenosine triphosphate (ATP) (3-6). In photosynthesis, PCET reactions power the chemically challenging light-driven oxidation of water (7-10), releasing dioxygen to our atmosphere, whilst various metalloenzymes rely on PCET reactions to catalyze central chemical conversion steps of the metabolism (11-13).

Ribonucleotide reductases (RNRs) are ancient enzymes, responsible for the *de novo* synthesis of deoxyribonucleotides from ribonucleotides that provide the building blocks for DNA in all cells (14, 15). All RNRs share a common strategy, converting a conserved cysteine residue into a transient thiyl radical (Cys•) that drives the reduction of ribonucleotides. (16) However, different subclasses of RNRs achieve this radical chemistry by distinct mechanisms. Class I RNRs comprise two subunits, the large R1 subunit, responsible for the ribonucleotide reduction, and the small R2 subunit, which generates and stores the radical used in the reduction process. The canonical class Ia R2s rely on a bimetallic iron core to activate and split dioxygen, generating a high-valent (Fe^III^/Fe^IV^) metal site, which in turn, creates a tyrosyl radical (Y•, *E*_m_ of *ca*. 800 mV vs. NHE) next to the metal center, first described more than 50 years ago (17). Y• subsequently drives long-range (>30 Å) radical transfer and PCET reactions (16, 18, 19), most likely *via* conserved tryptophan and tyrosine residues in R2. After crossing the protein interface, the radical is transferred to the R1 subunit, which results in the reactive Cys•, driving the ribonucleotide reduction (16). Upon completion of the reaction, the radical is regenerated and returned to the R2 subunit for reuse. The redox potentials of the radicals in R2 and R1 must thus be specifically tuned to allow for the reversible transfer. Yet, despite significant structural (20-24), biochemical (14), spectroscopic (25-28) and computational (29-34) insights over the last decades, especially on the class Ia RNR from *E. coli*, the mechanism of this reversible long-range PCET reaction remains elusive and much debated, in particular the principles and energetics of the reversible charge transfer process between the R1/R2 subunits. Despite the recently resolved structure of the *E. coli* R1-R2 holo-complex (21, 24), a structure of a non-perturbed subunit interface has not yet been determined. In this regard, previous studies suggest that the R1-R2 interface is dynamic, with distinct redox-driven conformational dynamics during the forward and reverse charge transfer (21, 35-41).

While the *E. coli* RNR drives the ribonucleotide reduction by employing a bimetallic transition metal core, nature has also evolved even more remarkable solutions to power the long-range PCET. In this regard, the recently discovered metal-free class Ie RNR, R2 (R2e), catalyzes the ribonucleotide reduction without the metal center. Instead, R2e relies on a post-translationally modified tyrosine residue, forming a 3,4-dihydroxyphenylalanine radical (DOPA•) (Fig. 1A, B) (22, 23, 42), which exhibits a unique stability, persisting for several days at room temperature. However, the metal-free R2e-RNRs impose a particular challenge for the reversible long-range charge transfer between R1 and R2, as the mid-point potentials of meta-hydroxylated phenols (catechols), forming the DOPA cofactor, are generally much lower (200-300 mV) relative to tyrosine radicals (14). This creates a thermodynamic challenge in R2e, as this process is already *ca*. 200 mV uphill in the canonical (class Ia) RNRs that rely on the redox-active tyrosine (see above) (43). We therefore expect that DOPA• is significantly redox-tuned in R2e to enable the reversible radical transport to the catalytic radical cysteine in R1.

**Fig. 1.**
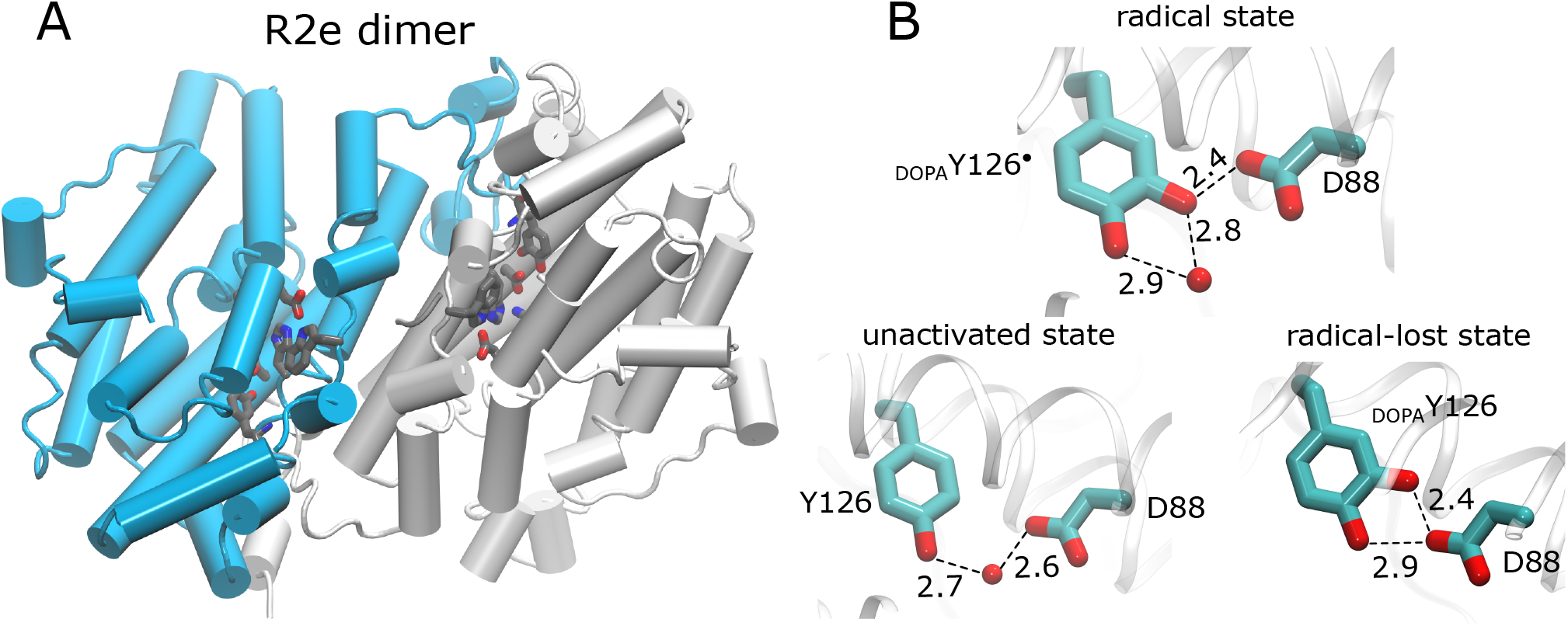
Structure of the R2e ribonucleotide reductase and its active site. (A) Structure of the R2e dimer. Two monomers are shown in blue and grey, respectively, with the location of the active site depicted in both monomers. (B) The protein surroundings around DOPA/tyrosine in the radical state, unactivated state, and the radical-lost (reduced) state, resolved using XFEL crystallography. Hydrogen-bonding (O…O) distances between Tyr126_DOPA_– Asp88 are shown in ångströms.

Femtosecond x-ray free-electron laser (XFEL) experiments in R2e of *Mesoplasma florum* demonstrated that the DOPA• forms an unusually short hydrogen bond with the adjacent conserved Asp88, with an O…O distance of 2.41 Å ± 0.05 Å (22) (Fig. 1B). Such a structure is indicative of a low-barrier hydrogen bond (LBHB) (44, 45), and could explain the unique stability of DOPA•. While LBHBs can be significantly stronger than normal hydrogen bonds (with donor/acceptor distance >2.5 Å) (46), their role in tuning the catalytic properties of enzymes remains controversial and poorly understood (47, 48). Yet, although short hydrogen bonds do not always result in barrierless proton transfer reactions or a catalytic benefit (49, 50), we hypothesize that the unique structure of DOPA• could provide the catalytic power of R2e.

To address this challenging question, we employ here a multidisciplinary approach by combing multiscale hybrid quantum/classical molecular dynamics (QM/MM-MD) and atomistic molecular simulations based on XFEL data with site-directed mutagenesis experiments, x-ray crystallography, and molecular spectroscopy. Our integrative theory-driven experimental approach provides a powerful methodology to probe the mechanistic principles of this long-range PCET process, allowing us to link together the energetics and dynamics of the DOPA• in R2e (*via* QM/MM, MD, and redox calculations), with distinct spectroscopic fingerprints (*via* EPR, ENDOR, FTIR, computational spectroscopy) and high-resolution structural data (XFEL, x-ray crystallography). Here, we study properties of the DOPA• within the isolated R2e system, and compare it with the same system, without the radical (*i*.*e*. “*unactivated*” state) or in the ‘radical lost’ state, obtained by chemically reducing the radical with a proton and an electron. Our combined findings suggest that an LBHB provides the underlying redox-tuning principles that enable the long-range radical transfer responsible for ribonucleotide reduction.

## Results and Discussion

### Low-barrier hydrogen bond results in a delocalized proton

The XFEL structure of the R2e radical state shows a short O…O bond distance of 2.41 Å between DOPA• and Asp88 that extends along a hydrogen-bonded network via Lys213, His122, Asp212 to Trp52, implicated in the radical transport (22). To understand the protonation dynamics in this network, we performed QM/MM-MD simulations on the R2e in its radical state, and compared these to simulations of the reduced (*i*.*e*. the ‘radical-lost’) state (see *Introduction*, SI Appendix, Fig. S1). To account for accurate polarization effects, we treated the entire hydrogen-bonding network and surrounding residues at the quantum mechanical level, while considering the biological surroundings on the classical level (see *Methods*, SI Appendix, Fig. S1).

The QM/MM-MD simulations indicate a barrierless proton hopping between DOPA• and Asp88 that results in a delocalized proton between the two residues (Fig. 2A, B). The QM/MM simulations show a short average O…O distance of 2.43 Å (Fig. 2A, B), while nuclear quantum effects (NQE) further reduce the O…O distance by 0.04 Å (Fig. 2B and SI Appendix, Table S1), consistent with the XFEL data. The LBHB is stabilized by coupled proton transfer reactions within the hydrogen-bonded network between Lys213/His122 and His122/Asp212 that resonate with the proton transfer at DOPA•/Asp88 (Fig. 2D and SI Appendix, Fig. S2-S4), with several energetically accessible charge-shifted microstates along the network (*i*.*e*., the proton shifted to Lys213, His122 and/or Asp212, see SI Appendix, Fig. S2). DOPA• strongly interacts with a dangling water molecule, forming an *out-of-plane* hydrogen bond with the para-oxygen (O_para_), which holds the majority of the spin density (SI Appendix, Fig. S3E, F). The LBHB character is supported by the flat free energy profile for the proton transfer between DOPA• and Asp88 (Fig. 2C and SI Appendix, Fig. S4), whilst *in silico* deuteration of the hydrogen-bonded network suggests that the proton delocalization is sensitive to isotope effects (SI Appendix, Figs. S4C, S21), with more transitions observed between DOPA and Asp88 in the protonated state relative to the deuterated simulations. As suggested by our structure optimizations (Fig. 2B), NQEs further decrease the DOPA– Asp88 bond distance by *ca*. 0.04 Å, and may thus contribute in forming the tight LBHB system, facilitating the proton delocalization.

**Fig. 2.**
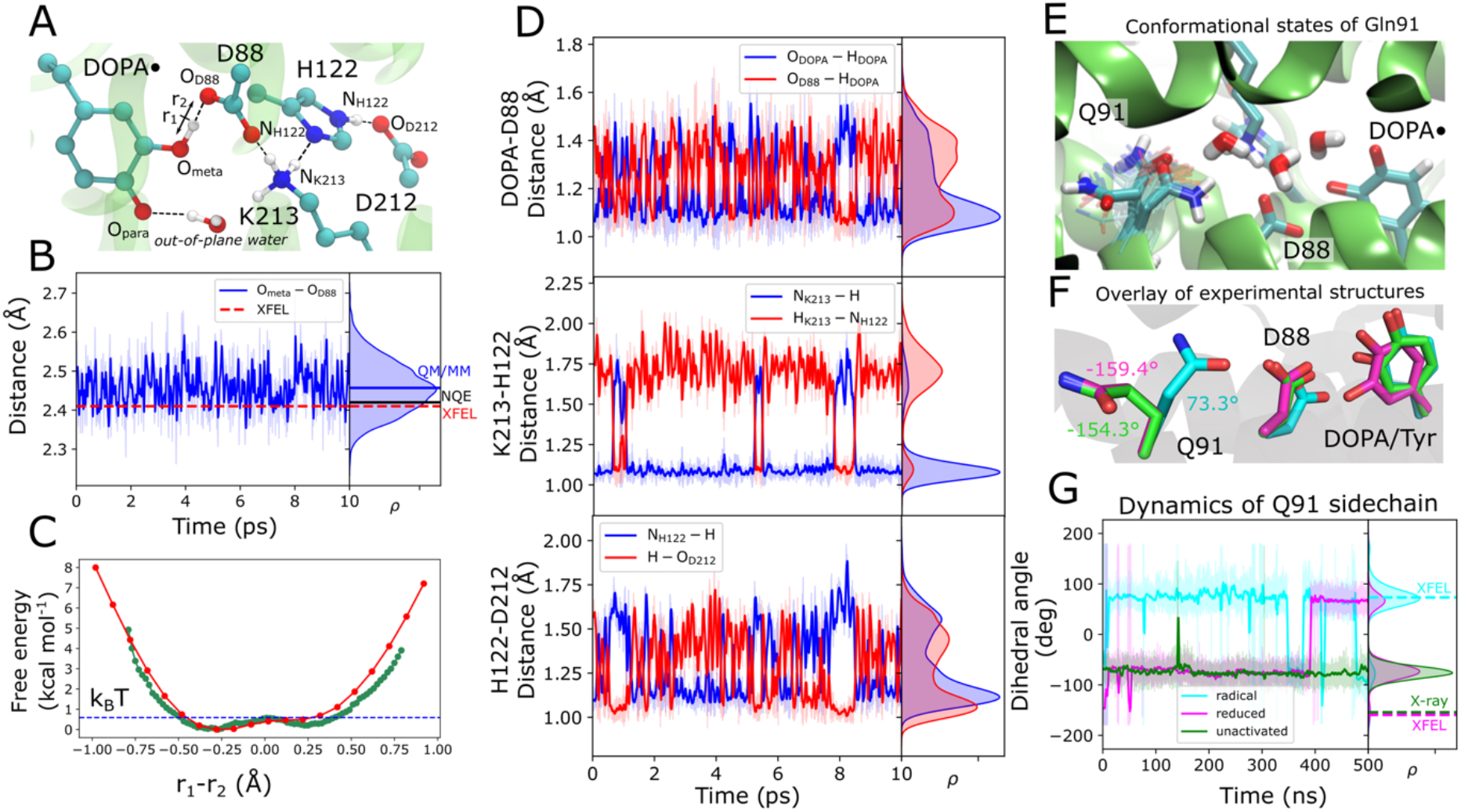
Structure, energetics, and dynamics of the DOPA• low-barrier hydrogen bond. (A) Hydrogen-bonded network extending from DOPA• through Asp88, Lys213, His122 to Asp212, located *ca*. 15 Å away. The key proximal *out-of-plane* water molecule adjacent to DOPA• is depicted. (B) Distance between DOPA• (O_meta_ atom) and Asp88 (O_D88_ atom) from QM/MM-MD simulations. The red dashed horizontal line represents the corresponding XFEL-derived distance. The simulation-averaged O–O distance and the nuclear quantum effects (NQE)-corrected O–O distance are also indicated (Supplementary Table 1). (C) Free energy profile for proton transfer between DOPA• and Asp88, obtained based on QM/MM free energy simulations (shown in green). *r*_1_ and *r*_2_ are DOPA•–H and H–Asp88 distances, respectively. The potential energy surface (PES) is overlaid in red. (D) Hydrogen-bond distances for the DOPA–Asp88, Lys213–His122, and His122–Asp212 from QM/MM-MD simulations. (E) Conformational dynamics of Gln91 observed during classical MD simulations of R2e in the presence of DOPA•. The conformational states of Gln91 influence the directionality of the water-mediated hydrogen-bond network. (F) Overlay of experimental structures of radical state (blue, PDB ID: 8bt3), radical-lost state (purple, PDB ID: 8bt4), and the unactivated state (green, PDB ID: 6gp3), showing conformational switching of Gln91, showing the dihedral angle (Cβ-Cγ-Cδ-Nε2) of Gln91 (in F-G). (G) Conformational dynamics of Gln91 for the radical, radical-lost, and unactivated states with relative populations of the respective dihedral −75º and +75º marked in the figure. The radical state samples also a minor population (<10%) an *inactive*-like state. The corresponding dashed lines indicate experimental values.

Our simulations suggest that the conformation of the proximal water molecule next to DOPA• is essential for modulating the properties of this LBHB network. When this water molecule is enforced into an *in-plane* orientation (SI Appendix, Fig. S2), the LBHB switches into a regular hydrogen bond (SI Appendix, Fig. S4B), with the proton localized on Asp88 (SI Appendix, Fig. S5), but with charge-shifted intermediates along the proton array that are energetically uphill by a few kcal mol^-1^ (SI Appendix, Fig. S2). In stark contrast to the radical state, the reduced (‘radical-lost’) DOPA shows a conformation, where the meta- and para-hydroxyl groups share hydrogen bonds with Asp88, resulting in O_meta_…O_D88_/O_para_…O_D88_ distances of 2.44 Å/2.85 Å, respectively (SI Appendix, Fig. S6). Despite the short distances, the proton remains bound to DOPA, and forms a regular hydrogen bond, involving a localized proton (SI Appendix, Fig. S7). We also identify an alternative minor conformation, where O_meta_ forms a hydrogen bond with Asp88, while the bridging water molecule is enforced into an *out-of-plane* orientation, also characterized by a regular hydrogen bond (SI Appendix, Fig. S8).

### Modifying the LBHB network results in loss of DOPA•

Atomistic MD simulations of the radical and reduced states on longer (microsecond) timescales result in the formation of a water array between DOPA and the outside bulk solution, which also link to conformational changes in the conserved Gln91 and Leu183 residues, as well as the water conformation next to DOPA (*in-plane* / *out-of-plane*). Our simulations suggest that the redox state could modulate the conformational ensemble (Fig. 2E and SI Appendix, Fig. S9), consistent with observations in the XFEL data (Fig. 2F, SI Appendix Fig. S9D, E, *cf*. also Ref. (22)). In this regard, the conformational flip of Gln91 stabilizes the hydrogen-bonding configuration of the DOPA sidechain underlying the LBHB arrangement (Fig. 2E-G). In contrast, when DOPA• is simulated with the *in-plane* water (extracted from MD simulations of the reduced state), the LBHB switches into a regular hydrogen bond, with the proton bound to DOPA• (SI Appendix, Fig. S8). These observations support that the configurations dominating the reduced state prevent proton delocalization along the extended hydrogen-bonded network (SI Appendix, Figs. S6-S8). The *in-plane* water orientation is locally stable on picosecond timescales of the QM/MM-MD simulations, but globally shifts towards the native-like radical conformation in the longer classical MD simulations, as also supported by the XFEL data (SI Appendix, Fig. S9D). Interestingly, mutation of Gln91 (Q91S, Q91L, see Fig. 2E) leads to a significant loss of DOPA• (Fig. 3A,B), as revealed by electron paramagnetic resonance (EPR) spectroscopy, suggesting that the residue could have a role in controlling the micro-solvation underlying the redox-tuning effects, or alternatively, perturb the interaction with the activating NrdI during the initial radical-formation process (23, 42). Moreover, substitution of Leu183 (L183A), a residue stabilizing the extended hydrogen-bonded network, also results in drastically reduced radical formation in EPR (Fig. 3A,B). In this regard, we observe a decrease in the hydration state of the cavity in the *in silico* variants L183A, Q91S, and Q91L, suggesting together with our EPR data that these residues are important for finetuning the hydration state around DOPA (SI Appendix, Fig. S23).

**Fig. 3.**
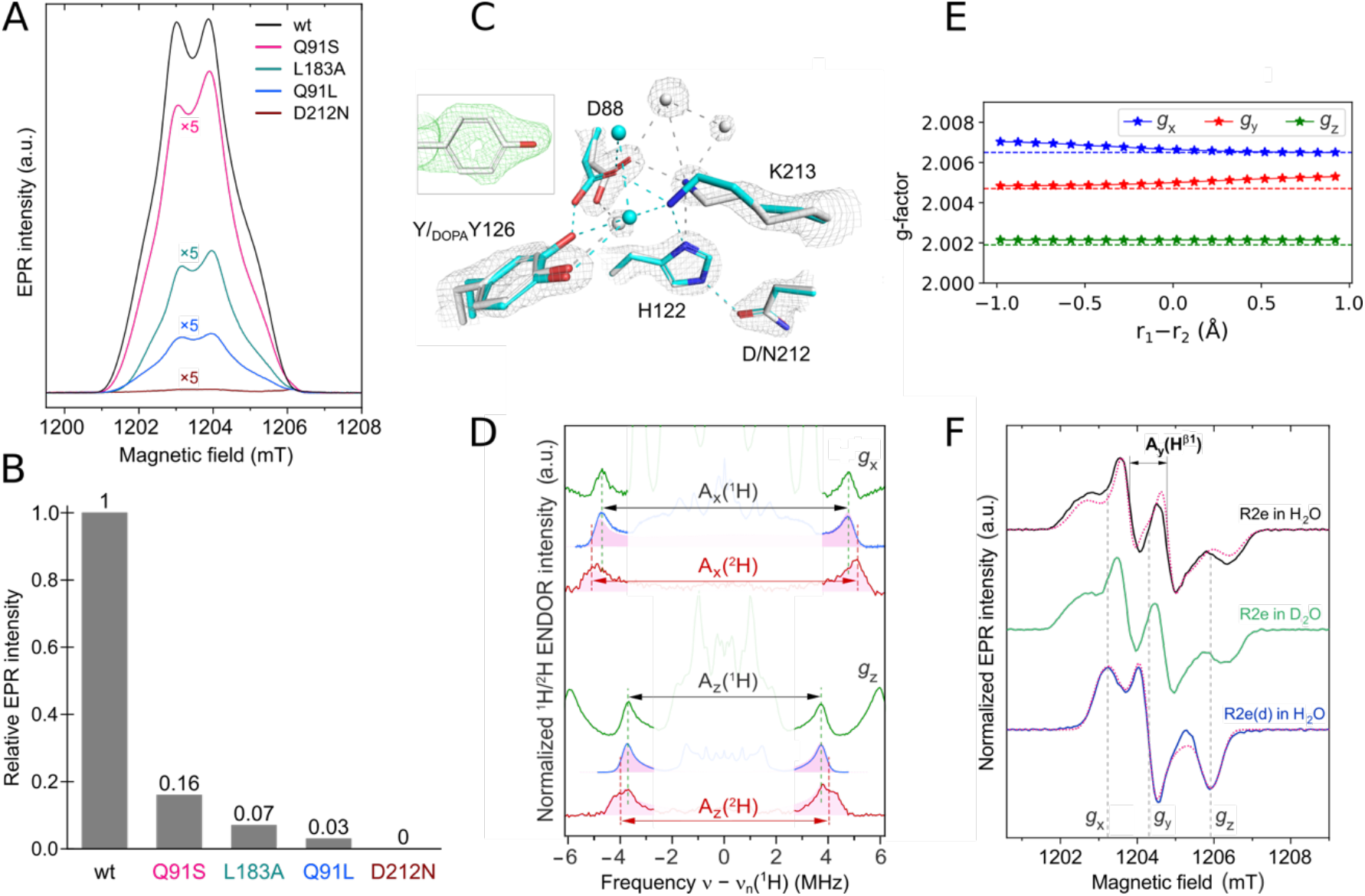
Directed mutagenesis and EPR characterization of DOPA•. (A) Q-band EPR spectra of WT-R2e containing DOPA• along with R2e variants, harboring substitutions either along the hydrogen-bonding network or within the DOPA hydrophobic cavity. (B) Relative intensities of the DOPA• EPR signal observed for WT and mutants. (C) Structural overlay of the active XFEL-determined WT-R2e structure containing the DOPA radical (*cyan*, PDB ID: 8BT3) with the structure of the D212N variant (*white*). The D212N mutation prevents the DOPA formation, stabilizing the inactive tyrosine state. The grey map is the 2F_o_-F_c_ map contoured at 2σ. *Inset:* omit map contoured at 1.2σ for Tyr126, confirming the absence of the DOPA modification. (D) Q-band Davies ENDOR spectra of the exchangeable LBHB in R2e, recorded along two canonical orientations of the DOPA• *g*-tensor (*g*_x_ and *g*_z_). The green and red traces correspond to protonated R2e in D_2_O: the green lines show the ^1^H ENDOR spectra, and the red lines show the corresponding ^2^H ENDOR spectra, scaled to the ^1^H gyromagnetic ratio. The blue traces show the ^1^H ENDOR spectra of perdeuterated R2e in H_2_O. The spectra were recorded at two canonical magnetic field positions corresponding to the *g*_x_and *g*_z_ orientations of the *g*-tensor. Vertical dashed lines highlight the difference in hyperfine couplings (A_x_, A_z_) between ^1^H and ^2^H for hydron between the LBHB. The pink-filled traces represent simulations for the LBHB. Signals from other protons are shown in pale colors to enhance the visibility of the LBHB contributions; see SI Appendix, Table S12 for the experimentally-fitted parameters and simulation of spectra. Signals from other protons are shown in pale colors to enhance the visibility of the LBHB contributions. (E) DFT-computed *g*-tensor of DOPA• along the proton-transfer coordinate. The colored dotted lines show experimental *g*-values. (F) Normalized pseudo-modulated EPR spectra of R2e in H_2_O (*black*) and D_2_O (*green*) buffers, and per-deuterated R2e in a H_2_O buffer (*blue*). Three turning points and the hyperfine splitting at *g*_y_ due to the Cβ-proton with A_iso_^β1^ ≈ 30 MHz are indicated with simulated spectra shown in pink.

To further probe how the extended hydrogen-bonded network affects the LBHB, we next introduced the D212N substitution, with the rationale of inverting the polarity of the hydrogen-bonded array (SI Appendix, Figs. S7B, S10-12). As expected, this mutation prevents the proton transfer between His122–Asn212 in our QM/MM-MD simulations, but surprisingly, also blocks the proton delocalization along the DOPA–Asp88–Lys213– His122 network (SI Appendix, Figs. S10, S11). To experimentally probe the consequences of these perturbations, we produced the D212N variant and determined its x-ray structure at 1.7 Å resolution (Fig. 3C and SI Appendix, Table S3). Remarkably, the D212N variant shows an unmodified tyrosine residue (Fig. 3C), as also supported by our EPR experiments (Fig. 3A,B), and suggesting that the extended hydrogen-bonded network could stabilize DOPA• and/or affect its initial formation. Taken together, these findings suggest that proton delocalization is important for the catalytic transformation of tyrosine into DOPA.

### Electronic and vibrational signatures of the LBHB and DOPA•

We next characterized the DOPA radical by EPR and electron nuclear double resonance (ENDOR) spectroscopy (Figs. 3A,B,D,F and SI Appendix, Figs. S13-S14) that provide powerful methods to study the electronic properties of radicals in proteins. To this end, we studied both the protonated as well as deuterated forms of R2e (see SI Appendix, *Methods*), and compared these to *g*-tensors and hyperfine couplings computed based on the QM/MM simulations (Fig. 3E and SI Appendix, Table S5-S8). Our EPR spectra confirm the formation of the DOPA radical, consistent with previous observations (23). Our QM/MM calculations revealed a rhombic *g*-tensor (*g*_x_>*g*_y_>*g*_z_), in excellent agreement with the experimental data, although the distinct water orientations, controlling the LBHB network, resulted in nearly identical *g*-tensors (Fig. 3E and SI Appendix, Fig. S5F). In this regard, the *g*_x_component, which is commonly sensitive to the local protein environment, remains invariant with respect to the position of the proton along the DOPA•–Asp88 hydrogen bond, as the majority of the spin density localizes on O_para_, and remains unchanged upon conformational switching (SI Appendix, Figs. S5E, S10E). Also in line with the QM/MM data underlying the LBHB state (Fig. 2 and SI Appendix, Figs. S3, S5), the ENDOR-detected hyperfine coupling of the Cβ hydrogen of DOPA•, which generally depends on the dihedral angle between the Cβ-H bond and the phenyl ring plane (51), supports the *out-of-plane* water configuration (SI Appendix, Figs. S13-S14). The ENDOR spectra show a stronger hyperfine coupling for deuteron between DOPA• and Asp88 than for the proton (normalized to the ratio of the gyromagnetic ratios), indicating that the former is located somewhat closer to DOPA• than the latter (Fig. 3D and SI Appendix, Figs. S4, S13-S14, Tables S5-S6, S12). At the same time, the dihedral angle is largely preserved upon deuteration of Cβ, as evidenced by ^1^H/^2^H ENDOR (SI Appendix, Table S12 and Fig. S13-S14), which implies that deuteration does not significantly perturb the structure of the DOPA radical overall. Thus, a pronounced isotope effect is only detected for the hydron between DOPA• and Asp88, in agreement with other reported systems with proposed LBHBs, where the isotope effect manifests itself as a change in the chemical shift (52, 53). Taken together, our EPR and ENDOR data (Fig. 3D, SI Appendix, Table S12 and Fig. S13-S14) further support the LBHB in the DOPA• state.

To probe the proton delocalization effects, we next analyzed vibrational signatures based on our QM/MM calculations. We find that the delocalized proton manifests in a unique vibrational mode linked to the DOPA•-O…H…O-Asp88 stretch in the high frequency (>2000 cm^-1^) range based on our QM/MM calculations (see SI Appendix, *Methods*). This vibration is highly sensitive to the O…O bond distance (Fig. 4A), and correlates with the formation of a local electric field at the moving proton, resulting from a vibrational Stark effect (VSE). In comparison to the reduced form, DOPA• shows characteristic vibrations appearing in the ranges of 1500–1800 cm^-1^, 2160 cm^-1^, and >3100 cm^-1^ (Figs. 4B and SI Appendix, Fig. S18-S19), with characteristic differences around 3100–3300 cm^-1^, 2600 cm^-1^, 2160 cm^-1^, and 1600–1700 cm^-1^ that are related to the LBHB in the DOPA• state. The vibrational difference spectrum computed on the basis of these models, shows vibrational features in the range of 1500–1800 cm^-1^; at 2160 cm^-1^; as well as unique features in the higher frequency range of 2600 cm^-1^ and 2900–3500 cm^-1^ that could arise from changes in the LBHB network upon reduction of the radical (Fig. 4B and SI Appendix, Fig. S18-S19).

**Fig. 4.**
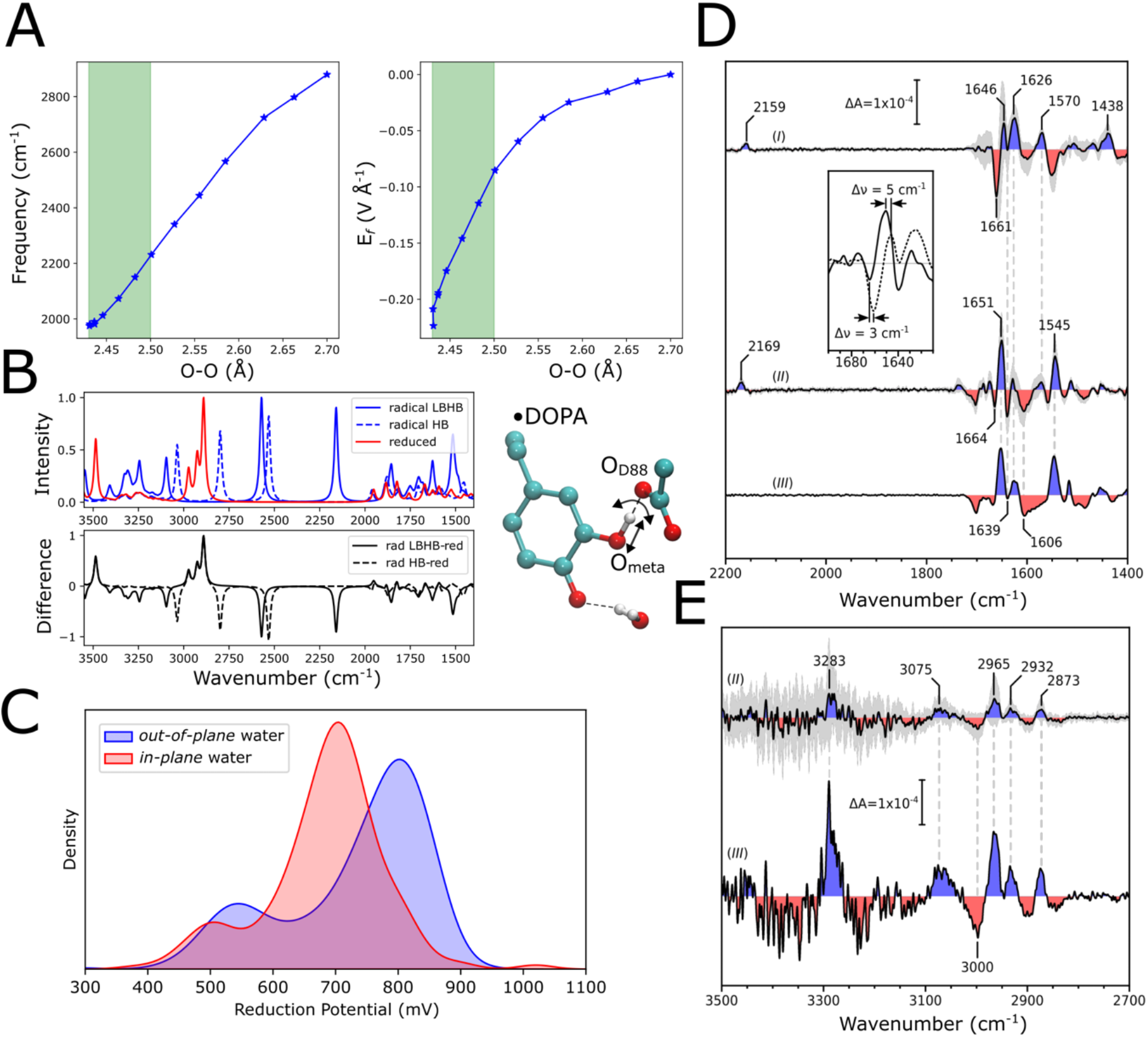
Redox properties and vibrational spectroscopy of the LBHB. (A) *Left*: Correlation between the O–O distance (DOPA•–Asp88) and the O–H stretching frequency, indicating characteristics of the proton delocalization. *Right*: Electric field magnitude along the proton transfer coordinate, computed at its midpoint, highlighting the influence of the protein environment on proton transfer energetics. (B) *Top*: Computed vibrational spectra from the LBHB configuration (with *out-of-plane* water, solid line) and a regular hydrogen bond (HB, *in-plane* water, dashed line). *Bottom*: difference spectra of reduced *minus* radical in the LBHB (solid) and HB (dashed) configurations (see *Methods*). *Left*: structure with characteristic vibration linked to the LBHB state (see Supplementary Fig. 18C, also for comparison to experimental spectra) (C) Reduction potential of DOPA• from QM/MM models with *in-plane* and *out*-*of*-*plane* water molecules, highlighting the role of water orientation in tuning redox properties. (D) Experimental FTIR difference spectra capturing vibrational signatures associated with the DOPA radical depletion and LBHB characteristics in the 2200–1400 cm^-1^ frequency range. FTIR difference spectra were recorded upon quenching of DOPA•-R2e, after addition of *I*) HU in D_2_O buffer, *II*) HU in H_2_O buffer, and *III*) NMHA in H_2_O buffer. *Inset*: Magnification of 1700–1610 cm^-1^ regime, showing spectral shifts upon H/D solvent exchange (solid line: H_2_O, dashed line: D_2_O). (E) High-frequency FTIR difference spectra recorded between 3500–2700 cm^-1^ upon quenching of DOPA•-R2e after addition of *II*) HU in H_2_O buffer, and *III*) NMHA in H_2_O buffer. Shaded grey areas indicate one standard deviation calculated between different experimental replicates, and vertical bars show the difference absorbance scale; relevant peak positions are labelled, with vertical dashed lines indicating bands appearing at identical frequencies.

To experimentally assess the vibrational features of the LBHB network resulting from the DOPA• state, we performed *in situ* Attenuated Total Reflection-Fourier Transformed Infrared (ATR-FTIR) spectroscopy, which provides a powerful methodology to address the vibrational fingerprint of proteins, and has also previously been used to study RNRs (54-57). Due to the many overlapping signals in the protein, the method relies on recording a difference spectrum, which we studied here by quenching the radical state with hydroxyurea (HU) or N-methylhydroxylamine (NMHA) (SI Appendix, Fig. S16). The resulting “*radical*-minus-*reduced*” FTIR difference spectra show consistent difference bands at 1665–1625 cm^-1^ upon formation of the ‘radical-lost’ state, regardless of the employed quencher (Fig. 4D). A negative/positive feature peaking at 1664^-^/1651^+^ cm^-1^ exhibits H/D sensitivity, and shifts to 1661^-^/1646^+^ cm^-1^ in D_2_O buffer, consistent with perturbed hydrogen-bonding and C=O decoupling in environments with exchangeable protons. Other characteristic spectral features include bands between 1560–1535 cm^-1^, which shift towards ∼1438 cm^-1^ in D_2_O. A unique positive band between 2170–2158 cm^-1^ is observed only upon quenching with HU, but is absent in NMHA-induced spectra, suggesting a specific HU-derived side reaction. Features in the higher frequency range (3300–2850 cm^-1^) emerge exclusively in H_2_O (Fig. 4E), likely corresponding to X-H stretching vibrations that may reflect reorganization within the LBHB network upon reduction of the radical (Fig. 4B). Our semi-quantitative comparison with the QM/MM-difference spectra (see Figure 4B, D, and SI Appendix, Fig. S18-S19), computed based on the harmonic approximation, suggests that several bands arise from the DOPA•-O–H–O-Asp88 vibrational modes and the C=O bond stretch, whereas the multiple shifts upon deuteration imply that the exchangeable protons are linked to the radical state (see *Methods*).

### Functional consequences of the LBHB for long-range radical transport

Under standard conditions, DOPA (catechol) has a lower oxidation potential by 200-300 mV relative to tyrosine, placing it in the 500-600 mV potential range. As discussed above, such potentials would render the radical transport to R1 highly endergonic by +0.3–0.4 eV, as the reaction in the canonical RNRs is already thermodynamically uphill by +0.1–+0.2 eV. To probe possible redox tuning effects in the metal-free R2e, we estimated redox potentials of DOPA based on our QM/MM trajectories using Poisson-Boltzmann electrostatic calculations coupled to Monte Carlo sampling of redox and protonation states (PBE/MC). To this end, we computed the intrinsic electron affinity based on quantum chemical calculations, followed by calculation of protein solvation effects within the PBE/MC framework (see *Extended Methods*). Although we lack experimental reference data for R2e, the PBE/MC method generally performs rather well in benchmarking calculations with an average error of *ca*. 60 mV for organic radicals relative to experimental data (58, 59). Our redox calculations suggest that the delocalized proton, together with the ensemble of energetically degenerate microstates along the hydrogen-bonded network, stabilize the radical, and result in a rather strong redox-tuning effect mainly by electrostatic interactions that upshifts the DOPA oxidation potential by >+300-400 mV relative to the normal hydrogen-bonded configuration, computed based on the radical lost state (Fig. 4C, SI Appendix, Fig. S17). These results suggest that the LBHB network could enable the isoenergetic radical transport process in R2e, and providing a basis for future electrochemical studies of redox tuning effects in this system. It is interesting to note that LBHBs have also previously been reported to result in large redox tuning effects (50), (60, 61), suggesting that the effects could generally reflect electrostatic tuning of protein function. It is also important to note that given the distinct cofactor composition and the overall architecture of class Ie versus class Ia RNRs, we do not assume an identical thermodynamic landscape in these systems. Our interpretations of redox tuning are therefore limited to relative trends within the metal-free (class Ie) systems studied here. However, the observation of the specific hydrogen-bonded network and redox-tuning in these metal-free RNRs, puts the DOPA radical on an even footing with the tyrosyl radical in the canonical RNRs, suggesting that the systems could have similar energetics. Taken together, our combined findings suggest the LBHB network results in strong redox-tuning effects that enable a reversible radical transport process that provides the catalytic power for the metal-free RNRs.

## Conclusions

We have shown here using an integrative computational-experimental approach that the unique stability of the DOPA radical in the metal-free RNR (R2e) arises from a low-barrier hydrogen bond (LBHB) network, with a delocalized proton. We find that a proximal water molecule could act as a molecular switch that enables long-range charge transfer within an extended (>10 Å) hydrogen-bonding network, with distant mutations resulting in the loss of radical formation as shown by our QM/MM simulations, high-resolution x-ray structure, as well as EPR and, in part, also vibrational spectroscopic data. Based on computed redox-properties, we further suggest that the LBHB exerts a strong redox-tuning effect of the DOPA• that provides a basis for the energetically uphill long-range radical transport in metal-free (class Ie) RNR. On a general level, our findings provide support for the direct involvement of LBHBs in protein chemistry and the importance of quantum effects in enzyme catalysis.

## Materials and Methods

Technical details of the QM/MM models and QM/MM free energy simulations, DFT models, computational spectroscopy, nuclear quantum effects, proton delocalization effects, redox calculations, vibrational spectroscopy (ATR-FTIR), EPR and ENDOR spectroscopy, mutagenesis experiments, and protein crystallography are available in the SI Appendix.

## Acknowledgments

A.S. acknowledges the EMBO Long-Term Fellowship (ALTF 952-2022) for support. The authors acknowledge financial support of this work by the Swedish Research Council (2025-04607 and 2020-04081 to V.R.I.K., 2021-03992 to M.H.), the German Research Foundation (DFG) under Germany’s Excellence Strategy – EXC 2033–390677874 – RESOLV (to M.K.), the DFG via the Collaborative Research Center (SFB1078, to J.H. and V.R.I.K.), by the European Research Council (HIGH-GEAR 724394 to M.H.), by the Knut and Alice Wallenberg Foundation (2019.0251 and 2024.0220 to V.R.I.K and 2017.0275 and 2019.0436 to M.H.), and the Göran Gustafsson Foundation for Research in Natural Sciences and Medicine (to V.R.I.K.). We acknowledge the National Academic Infrastructure for Supercomputing in Sweden (NAISS 2025/1-33, 2024/1-28, 2023/1-31) for computational resources. We thank the staff of Diamond light source, especially of beamline i04 for their help with the data collection (proposal mx29948). We acknowledge the Deuteration & Macromolecular Crystallization (DEMAX) team at European spallation source (ESS) and DEMAX project number 756075 for providing help and materials for the per-deuteration of R2e.

## Author Contributions

V.R.I.K. and M.H. conceived and led the study; A.S. and V.R.I.K. wrote the manuscript; A.S., A.P.G.H., and V.R.I.K. performed multiscale simulations; A.S, M.C.P, A.P.G.H., and V.R.I.K developed new methods; J.J., R.K., and V.S., performed mutagenesis studies; Y.K, and M.K. performed EPR and ENDOR experiments; F.B., and J.H. performed FTIR experiments; J.J., R.H., and V.S. performed protein production and crystallography; A.S., J.J., Y.K., R.K., F.B., V.S., H.L., A.P.G.H., J.H., M.K., M.H., and V.R.I.K. edited the manuscript; A.S., J.J., Y.K., R.K., F.B., V.S., H.L., M.C.P., A.P.G.H., J.H., M.K., M.H., and V.R.I.K processed and analyzed data.

## Methods

### Multiscale Simulations and Computational Spectroscopy

QM/MM simulations, DFT models, and atomistic molecular dynamics simulations were constructed for the WT-R2e and mutants based on the experimentally resolved x-ray structures of the radical state (PDB ID:8bt3), the radical-lost state (PDB ID:8bt4), or the unactivated state (PDB ID: 6gp3). The simulations were used to compute free energy profiles for proton transfer, estimate nuclear quantum effects, EPR and vibrational properties, as well as redox properties. The μs-dynamics of the systems were used to study hydration effects and protein fluctuations based on atomistic MD simulations. See SI Appendix, Extended Methods for a detailed technical description.

### R2e protein production, mutagenesis and crystallization

R2e mutants were generated using the WT-R2e plasmid using a Quick Change II site-directed mutagenesis kit (Agilent), following their protocol, with primer sequences shown in SI Appendix, Table S4. WT-R2e and R2e-mutants were expressed and purified as previously described (23). Per-deuterated R2e was expressed using a previously published protocol (62). The R2e-D212N variant was crystallized, following crystallization conditions applied to WT-R2e (23). X-ray data for R2e-D212N was collected at the beamline i04 at the Diamond Light Source (UK). The crystal structure was solved with molecular replacement using PDB ID: 6GP3 as a starting model and iteratively refined. See SI Appendix, Extended Methods for a detailed description of mutagenesis, protein expression and purification, deuteration of R2e, and x-ray crystallography.

### Infrared, EPR, and ENDOR spectroscopy

All infrared spectra of R2e were measured using a Tensor 27 FTIR spectrometer (Bruker, DE) equipped with a Mercury Cadmium Telluride (MCT) detector and a DuraSamplIRII ATR using a 3-reflection, ∼45° silicon ATR crystal (Smiths, UK), as described in Ref. (63). Reaction-induced difference spectra were measured by quenching the radical-active DOPA•-R2e with hydroxyurea (HU) or N-methylhydroxylamine (NMHA). The FTIR experiments were conducted at pH 7.0 (or pD 7.0) (25 mM HEPES or deuterated HEPES, 50 mM anhydrous NaCl, degassed) with the protein concentration of 1 mM - 1.6 mM under anaerobic conditions at ambient pressure and 25°C.

All EPR and ENDOR measurements on the DOPA•-R2e and variants were performed at the Q-band frequency (approx. 34 GHz) using 500–700 μM solutions of R2e in 1.6 mm quartz tubes. Pulse measurements were carried out at 65 K using a Bruker Elexsys E580 spectrometer equipped with a 150 W TWT amplifier, Bruker EN 5107D2 resonator, Oxford Instruments CF935 continuous-flow helium cryostat and Oxford Instruments MercuryiTC temperature controller. Q-band orientation-selective Davies ENDOR spectra were collected with stochastic detection at 65 K using an AR 600 W radiofrequency (RF) amplifier (AR 600A225A). See SI Appendix, Extended Methods for a detailed technical description of the spectroscopic methods.

Extended Materials and Methods, additional references, additional Tables and Figures can be found in the Supplementary Information. Computational models and force field parameters are available at zenodo: 10.5281/zenodo.15747346.

## Data availability

The Mf R2e-D212N mutant structure has been deposited in the Protein Data Bank with the following accession code: 9R5L. Computational models and spectral data are available at zenodo: 10.5281/zenodo.15747346.

