## Supplementary Information for "Low-barrier hydrogen-bond powers long-range radical transfer in the metal-free ribonucleotide reductase"

**This PDF file includes:**

**Extended Methods**

**Fig. S1** QM/MM, DFT cluster models, and atomistic MD simulations of R2e.
**Fig. S2** DFT models and energetics of the R2e radical state (S = 1/2).
**Fig. S3** QM/MM-MD simulations of WT-R2e in the radical state with the out-of-plane proximal water molecule.
**Fig. S4** Energetics of LBHB and H/D exchange.
**Fig. S5** QM/MM-MD simulations of R2e-WT in the radical state with the in-plane proximal water molecule.
**Fig. S6** QM/MM-MD simulations of the radical-lost state with two hydrogen-bonds between DOPA and Asp88 (Conf II).
**Fig. S7** DFT models and energetics of the radical-lost state.
**Fig. S8** QM/MM-MD simulations of the radical-lost state with one hydrogen-bonds between DOPA and Asp88 (Conf I).
**Fig. S9** Classical MD simulations of inactivated Y126 and DOPA in radical and radical-lost states.
**Fig. S10** QM/MM-MD simulations of R2e-D212N in the radical state with the out-of-plane proximal water molecule.
**Fig. S11** DFT models and energetics of the D212N variant of R2e in the radical state.
**Fig. S12** QM/MM-MD simulations of R2e-D212N in the radical-lost state with the in plane proximal water molecule.
**Fig. S13** Orientation selective Q-band $^1H/^2H$ ENDOR spectra for the WT R2e radical state in water buffer sample.
**Fig. S14** Nomenclature of the DOPA• ring and C$\beta$ protons.
**Fig. S15** Quenching of radical-active DOPA•-R2e monitored through UV/Vis spectroscopy.
**Fig. S16** ATR-FTIR acquisition protocol and baseline correction procedure for reaction-induced difference spectra.
**Fig. S17** Redox properties of DOPA•.

**Fig. S18** Electric field and vibrational effects of the LBHB based on DFT models and H/D isotope effects.
**Fig. S19** Selected vibrational normal modes linked to DOPA/Asp88/proximal water.
**Fig. S20** Benchmarking the quantum chemical methods.
**Fig. S21** H/D isotope effects from QM/MM-MD simulations.
**Fig. S22** Analysis of MD simulations.
**Fig. S23** MD simulations of the L183A, Q91S, and Q91L variants, and the WT R2e in the radical state.

**Table S1** Nuclear quantum effects in R2e.
**Table S2** Benchmarking the molecular energetics.
**Table S3** X-ray data collection of R2e-D212N.
**Table S4** Primers for R2e mutagenesis.
**Table S5** Calculated g-tensors of DOPA•.
**Table S6** DFT computed hyperfine coupling constants for the out-of-plane water orientation.
**Table S7** DFT computed hyperfine coupling constants for the in-plane water orientation.
**Table S8** Variation of the DOPA• g-tensor and 1H hyperfine coupling constants along the LBHB proton transfer coordinate.
**Table S9** Modeled QM region in the QM/MM simulations.
**Table S10** List of QM/MM-MD simulations of various redox state of R2e.
**Table S11** List of classical MD simulations of various redox state of R2e.
**Table S12** Experimentally fitted hyperfine parameters for simulation of the ENDOR spectra.
**Table S13** List of protonation states used in MD simulations.

**Movie S1** LBHB network dynamics in DOPA•.
**Movie S2** Hydrogen-bonding network dynamics in the radical-lost state.

**SI References**

#### Extended Methods

#### Materials and Methods

##### 1. DFT models and methods benchmarking

Quantum chemical cluster models of the WT and D212N variant of the *Mesoplasma florum* R2e were constructed to probe the relative energetics of the different protonation arrangements in the hydrogen-bonded network of R2e. The models were constructed based on the experimental structures, where $Y_{DOPA}$126 is present in the reduced ("radical-lost", PDB ID: 8bt4)(1) or oxidized ("radical", PDB ID: 8bt3) states(1). For the reduced state model, Phe81, Leu84, Thr85, Asp88, His122, Ser125, $Y_{DOPA}$126, Leu183, Phe187, Ile206, Ile209, Asp212, and Lys213 were included in the DFT model. A similar DFT model was also created for the oxidized state of $Y_{DOPA}$126, but also including Gln91 and three resolved water molecules proximal to DOPA. The residues with a truncated backbone were terminated by a methyl group ($-CH_3$) at the $C_\alpha$ atom. The $C_\alpha$ atoms and the linker hydrogen atoms were fixed at their x-ray positions during the geometry optimization. The model comprised 180 atoms and 189 atoms for the "radical-lost" and "radical" states, respectively. All geometry optimizations were performed at the DFT level using B3LYP-D3/def2-SVP, with single point energy calculations performed at the ωB97M-V/def2-TZVPP(2, 3) level using an implicit solvation model (COSMO) with the dielectric constant set to 4.0, along with empirical D3 dispersion correction(4) and the m4 DFT integration grid.

For benchmarking energetics, single-point energy calculations were performed on the optimized structures using various DFT functionals (TPSS(5), TPPSh(6), B3LYP(7), M06-2X, B2PLYP(8), ωB97X-D(9) and ωB97M-V[2]) on the "radical" state models, comprising a smaller cluster with $Y_{DOPA}$126, Asp88, Lys213, and two water molecules hydrogen-bonding with $Y_{DOPA}$126 (total 56 atoms) to reduce the computational cost. The energetics was benchmarked against correlated Random Phase Approximation (RPA)(10) and domain-based local pair natural orbital coupled-cluster theory (DLPNO-CCSD)(11). All single point calculations were performed using correlation-consistent cc-pVTZ basis sets. Dispersion effects were included by the empirical D3 dispersion correction, together with a higher DFT integration grid (m4) and tighter SCF (scfconv 8). Implicit solvation effects were introduced via the COSMO scheme with the dielectric constant set to 4.0. All DFT and RPA calculations were performed using TURBOMOLE v7.5-7.7(12), and DLPNO-CCSD calculations(11) were performed using ORCA v6.0(13).

##### 2. System setup and atomistic molecular dynamics simulations

Classical molecular dynamics (MD) simulations were used to probe the conformational dynamics of $Y_{DOPA}$126 and its extended hydrogen bonding network. The simulations were performed with $Y_{DOPA}$126 in the radical-lost form (PDB ID: 8bt4), the radical form (PDB ID: 8bt3) and the unactivated form (PDB ID: 6gp3)(1, 14). The models were constructed based on the experimentally determined x-ray and XFEL structures of MfR2. Hydrogen atoms were added by assuming a physiological pH of 7.5, and missing residues were modeled using PyMOL. Protonation states of titratable residues were assessed using the H++ server as well as by performing Poisson-Boltzmann electrostatic calculations coupled with Monte Carlo (PBE/MC) (see section ***8. Redox potentials of DOPA***). The protonation states used in the simulations are listed in Table S13. The protein models were embedded in a TIP3P water box along with 150 mM NaCl. Parameters for the protein, $Y_{DOPA}$126, water and ions were based on the CHARMM36 force field(15). The total system size was *ca*. 150,000 atoms.

The MD setups were first energy minimized for 2000 steps, while restraining the heavy backbone atoms, followed by simulations in the *NVT* ensemble, with the heavy backbone atoms restrained. The system was then equilibrated for 250 ps at 310 K, controlled using Langevin dynamics. Subsequently, we conducted *NPT* equilibration for 2 ns, while maintaining the restraints on non-hydrogen backbone atoms. The pressure was kept at 1 bar, and the temperature was set to 310 K using Langevin dynamics. After the equilibration phase, all restraints were fully released, and an additional 5 ns unrestrained simulation was performed. The final snapshot from this trajectory was used as the starting structure for the production simulations, performed in an *NPT* ensemble with *T* = 310 K using the Langevin thermostat with a friction coefficient of 1 $ps^{-1}$, and pressure of 1 atm maintained using Nosé–Hoover–Langevin piston pressure control, and an integration timestep of 2 fs. Long-range electrostatic interactions were modeled using the Particle Mesh Ewald (PME) approach with a grid size of 1 Å. Production simulations were performed for each redox state of $Y_{DOPA}$126 and different permutations of protonation states along the $Y_{DOPA}$126-Asp88-Lys213-His122-Asp212 network. All MD simulations were propagated for 500 ns in duplicates (cumulative 3 μs). The MD simulations were performed using NAMD3.0(16). Trajectories were analyzed using VMD(17) and MDAnalysis(18). See Table S10 for a complete list of atomistic MD simulations, Table S13 for *protonation states used in the simulations,* Figure S22 for analysis of the convergence of MD simulations, and for comparison to experimental B-factors.

##### 3. QM/MM models and free energy simulations

For generating snapshots used in the QM/MM simulations, we repeated the minimization (see above), as well as the *NVT* and *NPT* equilibration, but by applying constraints to all heavy atoms within 8 Å of the central DOPA hydrogen-bonding network in monomer A. This protocol allowed the surrounding protein environment to relax while preserving the positions of key residues as resolved in the XFEL structure.

QM/MM-MD simulations were performed to probe the redox-driven dynamics of $Y_{DOPA}$126 and its central hydrogen bonding network extending from $Y_{DOPA}$126 to Asp212. For this purpose, QM/MM-MD simulations were carried out on the R2e dimer. All protein backbone atoms and residues (except hydrogens) 14 Å around the Lys213 were fixed to their experimentally derived coordinates. The last structural snapshot extracted from the classical simulations was used as an input structure for the QM/MM simulations. A complete list of the QM/MM-MD simulations and the description of the QM region are provided in Table S9 and Table S8, respectively. The QM region was treated at the B3LYP-D3/def2-SVP level of theory, while the MM system was modeled using the CHARMM36 force field. Covalent bonds along the QM-MM interface were cut at the $C_\alpha$-$C_\beta$ bond using the link-atom approach. The MM region within 14 Å of the QM region was flexible during the QM/MM simulations, whereas the remaining part was fixed. QM/MM simulations were performed using an integration timestep of 1 fs propagated at *T*=310 K, controlled using the Nose-Hoover thermostat. All QM/MM MD simulations were cumulatively propagated for 70 ps, with the entire QM/MM setup comprising around 83,450 atoms.

To understand the energetics of the LBHB between DOPA and Asp88 in the radical state, we optimized the QM/MM potential energy profile and performed QM/MM free energy simulations (see below). For optimizations of the potential energy surface, the proton transfer reaction was studied in models with both *in-plane* and *out-of-plane* water molecules. The QM region in these calculations was the same as in the QM/MM-MD simulations. The potential energy surface was studied by restrained structure optimization, with harmonic restraint of 3000 kcal $mol^{-1}$ $Å^{-2}$ placed on the reaction coordinate $R$, defined as a linear combination of bond forming ($r_1$) and bond breaking ($r_2$) distances within the hydrogen bond between the DOPA and Asp88, $R=r_1-r_2$ (Fig. S4). The high force constants were used here to effectively fix the positions of the reaction coordinate to a given value during structure optimizations, but implemented via a harmonic restraint due to technical aspects. To enhance the convergence, $R$ was scanned in both forward and backward directions until the converged profiles were obtained (defined by energy change <0.006 kcal $mol^{-1}$ in consecutive steps). The calculations were formed using B3LYP-D3/def2-SVP level of theory with the QM/MM calculations performed using the CHARMM-TURBOMOLE interface (19). Isotope effects of the radical state were studied by performing QM/MM-MD simulations with deuterium substitution (Fig. S21).

QM/MM free energy simulations of the proton transfer reaction were studied using umbrella sampling (US). The initial structures for US simulations were derived from the potential energy surface calculations (see above). The US windows were placed at 0.1 Å separation, resulting in a total of 16 windows, sampled using a harmonic restraint of 100 kcal $mol^{-1}$ $Å^{-2}$ placed on the proton transfer reaction coordinate $R$. Each US window was propagated for *ca.* 2.5 ps at $T$=310 $K$ (Nose-Hoover thermostat), with a timestep of 1 fs, resulting in cumulative QM/MM-MD sampling of 40 ps for each state. The free energy profiles were derived from the weighted histogram analysis method (WHAM, ref (20)) at 310 K using a convergence threshold of 0.0001 kcal $mol^{-1}$, and by estimating statistical errors based on bootstrap analysis (see Figure S4E for convergence). The QM region was the same as in the corresponding QM/MM-MD simulations, and treated at the B3LYP-D3/def2-SVP level.

#### 4. Computational EPR spectroscopy

EPR parameters (*g*-tensors and hyperfine coupling constant) for the $Y_{DOPA}$126 radical state were computed based on the optimized QM cluster models. The EPR parameters were computed using the TPSSh functional, which has provided accurate results for the tyrosyl radicals in Photosystem II(21) and class Ia RNR(22). The calculations were performed using the EPR-II basis sets, specifically tailored for the EPR calculations. The calculations were performed using the tight SCF criteria and a higher DFT integration grid (VeryTightSCF and DEFGRID3 in ORCA). The gauge origin for the g-tensor calculation was defined at the origin of the $Y_{DOPA}$126 ring. The *g*-tensors were computed using the coupled-perturbed self-consistent field approach (CP-SCF)(23) along with spin-orbit mean-field approximation to the Breit-Pauli operator(24) to treat spin-orbit coupling. The hyperfine coupling constants were computed for all protons on the $Y_{DOPA}$126 radical and for the proton directly hydrogen bonded to it ($H_2O$). All computations were performed using ORCA5.0(13)

#### 5. Nuclear quantum effects

To probe the role of nuclear quantum effects on the LBHB, geometry optimizations were performed using the nuclear–electronic orbital (NEO) method(25, 26), which treats protons quantum mechanically, *i.e.*, with a nuclear-electronic wavefunction. To this end, a cluster model consisting of $Y_{DOPA}$126, Asp88, Lys213, and His122 (46 atoms) with electron exchange and correlation treated at the B3LYP level, while the electron-proton correlation was treated with the epc17-2(27) functional. The electrons were treated using the def2-SVP electronic basis sets and the PB4-F2 protonic basis set(28), with all geometry optimizations carried out in the gas phase. The computations were performed using Q-Chem 6.0(29).

**6. Analysis of resonance effects in the LBHB network**

Sequential hydrogen bonding distances along the LBHB network were extracted from QM/MM-MD simulations (Fig. S4E). The time-dependent distances were transformed by fast Fourier transformation (FFT) into the frequency domain using the scipy.fft module in Python (30). The cross-correlation between distance timeseries $x_i$ and $y_j$ with $0 \leq i, j \leq N$ was computed according to,

$$z_k = \sum_{l=0}^{N-1} x_l \cdot y_{l-k+N-1} \tag{1}$$

with the scipy.signal module(30) to identify resonance effects/lag times along the LBHB network.

**7. Proton wave function along the LBHB network**

The 1D Schrödinger equation (1D-SE) was solved numerically for the proton moving on the potential energy surface extracted for the DOPA radical state. The 1D-SE was solved using the finite difference method. The proton wavefunction was described using an orthonormal finite basis set $\{\phi_i, i = 1, \ldots, N = 2000,$

$$\phi_i(x) = \begin{cases} \frac{1}{\sqrt{\Delta x}}, & if\ x_0 + \Delta x \cdot \left(i - \frac{1}{2}\right) < x \leq x_0 + \Delta x \cdot (i + \frac{1}{2}) \\ 0, & \text{else} \end{cases} \tag{2}$$

where $x_0 = -0.5$ Å and $\Delta x = \frac{\Delta X}{N} = \frac{1\,\text{Å}}{2000}$. The potential operator $\hat{V}$was approximated by a 12th degree polynomial *V*(*x*) fitted to the potential energy surface. The matrix representing the operator is a diagonal matrix, $V_{ii} = V(x_0 + \Delta x \cdot i)$. The kinetic operator $\hat{T}$ was approximated by the Laplacian matrix $T_{ij} = \frac{1}{\Delta x^2} \frac{\hbar^2}{2m} \Delta_{ij}$, where,

$$\Delta_{ij} = \begin{cases} 2, if\ i = j \\ -1, if\ |i - j| = 1 \\ 0, \text{else.} \end{cases} \tag{3}$$

Due to its sparse nature, the system of linear equations was solved using SciPy's "eigs" function, and implemented in Python.

#### 8. Redox potentials of DOPA

Redox potentials of the DOPA radical were calculated based on the QM/MM-MD trajectories of both the *in-plane* and *out-of-plane* water configurations. The redox potentials were calculated by solving the linearized Poisson-Boltzmann equation and by Monte Carlo sampling (PBE/MC)(31). To this end, the reference redox potential for the model DOPA compound in water was obtained from transferring the radical ($DOPA^{\bullet}$) and reduced ($DOPA^{-}$) from gas phase (*g*) to an aqueous solvent medium,

$$\Delta\Delta G_{solv}(\mathrm{DOPA}^{\bullet}/\mathrm{DOPA}^{-}) = \Delta G_{solv}(\mathrm{DOPA}^{-}) - \Delta G_{solv}(\mathrm{DOPA} \bullet\cdot) \quad (4)$$

where the solvation free energy for species X (X=$DOPA^{\bullet}$, $DOPA^{-}$) was computed based on,

$$\Delta G_{solv}(\mathrm{X}) = G_{\mathrm{wat}} - G_{\mathrm{g}}^{298K} \quad (5)$$

with the gas phase reduction potential defined as,

$$G_g^{298K} = E_o + ZPE + G_{TRV}^{298K}, \quad (6)$$

where $E_o$ is the electronic energy at the B3LYP-D3/6-31G(d,p) level, ZPE is the zero-point energy correction estimated from the molecular Hessian, and $G_{\mathrm{TRV}}$ are the translational, rotational, and vibrational energy contributions(32). The redox potential, $E_{redox}^{o}$ was computed based on,

$$n\,F\,E_{\mathrm{redox}}^{o} = \Delta G_g - \Delta\Delta G_{solv} + \Delta G_{NHE} \quad (7)$$

where *F* is the Faraday constant, 23.06 kcal $mol^{-1}$ $V^{-1}$, and $\Delta G_{NHE}$ = -4.43 eV is the normal hydrogen electrode (NHE). The solvation free energies were calculated based on the B3LYP-D3/6-311(d,p) optimized models of $DOPA^{\bullet}$ and $DOPA^{-}$, followed by fitting the electronic structure to RESP charges(33) and solvation radii(34), and embedding these into a polarizable medium (with an ε=80) using the PBE model. The redox potentials of DOPA in R2e,

$$E_{\mathrm{redox,R2e}}^{o} = E_{\mathrm{redox}}^{o} - \Delta\Delta G_{\mathrm{protein}} \quad (8)$$

was obtained from the PBE/MC sampling of 1000 structures per state (*in-plane* and *out-of-plane*) extracted from the QM/MM-MD simulations, based on the reference redox potential of the model compound (see above). For the protein solvation ($\Delta\Delta G_{protein}$), the $2^N$ protonation states of R2e were sampled using Metropolis Monte Carlo, allowing for single, double, and triple changes in the *N* protonation states of histidine (δ, ε, and δ+ε protons), tyrosine, glutamate (OE1 and OE2), aspartate (OD1, OD2 protons), lysine, and cysteine residues. The PBE/MC calculations were also used to assess the $pK_a$ of titratable residues in the active site of R2e (see Table S13). The quantum chemical calculations were performed using Q-Chem 6.0(29), the PBE equations were solved as implemented in the adaptive Poisson-Boltzmann solver (ABPS)(35), with the MC sampling performed using Karlsberg+(36).

#### 9. Electron affinity of DOPA

The electron affinity (EA) of DOPA was estimated within the QM/MM framework, based on

$$\mathrm{EA} = E(N) - E(N+1) \quad (9)$$

where E($N$) is the total QM/MM energy of the $N$-electron system and DOPA is in the radical state. The EAs were computed for snapshots extracted every 1 fs along the 10 ps QM/MM MD trajectory (10,000 structures). B3LYP/def2-SVP level of theory was used throughout and the protein effect was included via point charges.

**10. Vibrational properties**

Vibrational spectra of the DOPA radical state in the *out-of-plane* ("LBHB") and *in-plane* ("normal hydrogen bond") states, as well as in the reduced state, were computed based on the molecular Hessian estimated numerically from the optimized DFT cluster models (see *Methods*, Section 1) at the B3LYP D3/def2-SVP/ε=4 level. H/D isotope effects were computed by exchanging the mass of all hydrogen atoms to deuterium (2.01410200 a.u.), followed by re-calculation of the Hessian. The vibrational spectra were fitted to Lorentzian functions for computation of difference vibrational spectra, with the spectra empirically shifted to match a characteristic experimental (LBHB-DOPA) vibration at 2159 $cm^{-1}$. For analysis of vibrational Stark effects, vibrational spectra were computed by extracting the central DOPA radical, the proximal (*out-of-plane*) water molecule, Asp88, and Lys213, followed by restrained geometry optimization of the ($DOPA^{\bullet}$)-O–O-(Asp) distance from 2.43 to 2.70 Å (Fig. S19A), with the electric field computed at the moving hydrogen nucleus. All vibrational spectra were calculated using TURBOMOLE v7.5-7.7, with the vibrational normal modes analyzed using Molden v.7.3 (37) and JMol v. 16.2.

**11. Mutagenesis, protein expression and purification of R2e**

The pET28MfnrdFI plasmid (14) was used to generate the R2e mutants using a Quick Change II site-directed mutagenesis kit (Agilent), following their protocol, with primer sequences shown in Table S4. The successful mutation was confirmed by sequencing the plasmid. As the WT plasmid, the mutants contain an N-terminal, TEV-cleavable His-tag on the R2e protein for purification and an untagged WT MfNrdI gene that is co-expressed (see Ref.(14) for details). The mutants were expressed and purified similarly to the previously published protocol (14). Briefly, TB medium (Formedium) was supplemented with 50 µg $ml^{-1}$ kanamycin and 0.5% v/v of a starter culture containing *E. coli* BL21 DE3 cells transformed for each variant. The bacteria were grown in a LEX bioreactor (Epiphyte3) to an OD of 1.5, with the protein production induced by addition of 0.5 mM IPTG. The cells were incubated overnight at room temperature, harvested the next day, flash frozen and stored at -20°C until purification. For protein purification, the cell pellet was homogenized in lysis buffer (25 mM HEPES pH 7.0, 20 mM imidazole, 300 mM NaCl) and lysed by sonication. The lysate was cleared via centrifugation for 45 min at 40,000 g and 4°C and applied to a Ni-NTA column equilibrated in lysis buffer. After extensive washing, the protein was eluted from the column with elution buffer (25 mM HEPES pH 7.0, 250 mM imidazole, 300 mM NaCl), concentrated and injected on a Superdex S200 16/600 (Cytiva) column equilibrated in SEC buffer (25 mM HEPES 7.0, 50 mM NaCl). Relevant fractions were pooled, and the His-tag was cleaved of with TEV protease overnight at 4°C. Protease and potentially uncleaved R2e protein were removed by a reverse Ni-NTA step. The protein was concentrated to 20-25 mg $mL^{-1}$, aliquoted and flash frozen. Perdeuterated R2e was expressed using a previously published protocol(38). All the components were prepared in 99.8% deuterium oxide provided by the Deuteration & Macromolecular Crystallization (DEMAX) team at European spallation source (ESS). The WT MfNrdI plasmid containing *E. coli* was adapted to grow in deuterated minimal medium and induced by addition of 0.5 mM IPTG, cells were allowed to express for 16 hours before being harvested. Perdeuterated protein was purified the same way as WT R2e except in the last step it was buffer exchanged with deuterated SEC buffer.

**12. Crystallization, data collection and structure determination of R2e-D212N**

The R2e-D212N variant was crystallized, following crystallization conditions applied to WT-R2e(14). In this regard, a sitting drop vapour diffusion plate was manually set up with crystallization conditions ranging from 100 mM ammonium sulfate, 12-16% PEG 3350, and 100-200 mM calcium acetate and with a protein concentration of 20-25 mg $mL^{-1}$. 1 µL of protein was mixed with 1 µL of crystallization condition, and 0.2 µL of seed stock, which was produced from older crystals of the WT-R2e, and stored at RT. Crystals appeared after 1-2 days and were fished using 25% glycerol as cryoprotectant.

X-ray data was collected at the beamline i04 at the Diamond Light Source (UK). Data was collected at 100 K with a wavelength of 0.9537 Å. The data was reduced with XDS(39). The mtz file was tested with Xtriage of the Phenix Suite(40) and showed moderate anisotropy with lower resolution along the h axis. The crystal structure was solved with PHASER(41) using PDB ID: 6GP3 as a starting model. The model was built in Coot(42) and refinement was conducted with phenix.refine(43). Refinement included isotropic B-factors, occupancy, TLS parameters, and reciprocal space refinement. Water molecules were initially added with phenix.refine, manually adjusted and after an initial round of refinement manually edited. The final refinement was validated with MolProbity(44). A composite omit map using simulated annealing was generated with Phenix.

**13. Infrared spectroscopy**

***ATR-FTIR spectroscopy***

All infrared spectra were measured using a Tensor 27 FTIR spectrometer (Bruker, DE) equipped with a Mercury Cadmium Telluride (MCT) detector and a DuraSamplIRII ATR using a 3-reflection, ~45° silicon ATR crystal (Smiths, UK) essentially as described in ref (45, 46). The spectrometer was contained in an anaerobic chamber (Coy Laboratory Products, USA) constantly purged with $N_2$ gas. FTIR absorbance spectra were referenced to the blank ATR crystal and measured with 2 $cm^{-1}$ resolution between 4000–400 $cm^{-1}$. The background noise was quantified at $1x10^{-5}$ (root mean square) in the frequency regime between 2200–1400 $cm^{-1}$ by repeating blank measurements.

Radical-active DOPA•-R2e was solubilized in pH 7.0 or pD 7.0 buffer (25 mM HEPES or deuterated HEPES, 50 mM anhydrous NaCl, degassed) and used for the FTIR measurements. The protein concentration for experiments performed in $H_2O$ varied between 1 mM and 1.6 mM between several replicates, while the protein concentration in $D_2O$ was ~1.6 mM. The liquid sample was contained in a capped, custom 3D-printed small-volume liquid sample holder. The sample holder was made of polyvinylidene fluoride, commercially available as FluorX filament (3DXTECH, USA). All experiments were performed under anaerobic conditions at ambient pressure and temperature between 25°C and 26°C.

***Reaction-induced difference spectra***

Hydroxyurea (HU) and N-methylhydroxylamine (NMHA) were purchased from Sigma-Aldrich (USA) and solubilized in $H_2O$ or, in the case of HU, also in $D_2O$. Stock solutions of the reactants used were prepared at a concentration of 500 mM and stored in a -80°C freezer prior to experiments. Approximately 8 µL of radical-active DOPA•-R2e were added to a 200 µL Eppendorf tube together with 1/10 of the chemical quencher stock solution under an anaerobic atmosphere, resulting in a final quencher concentration of approximately 50 mM.

Immediately after mixing the protein and quencher, the liquid mixture was injected into the sample holder and absorbance spectra were measured every 30 s for a total time of 45 min. Absorbance spectra were averaged into six "classes" corresponding to ordered intervals of 5 min duration. These "classes" were then used to calculate difference spectra relating the later phases of the reaction to the earlier ones, namely by subtracting the classes corresponding to 0-5 min, 10-15 min, and 20-25 min from the classes averaged between 5-10 min, 15-20 min, and 25-30 min.

The quenching experiment with HU was repeated 5 times using $H_2O$ and 4 times for the $D_2O$ data. Difference spectra were calculated for each experiment as described above. This resulted in a total of 15 spectra for the $H_2O$ data and 12 spectra for the $D_2O$ data. The difference spectra in the low-frequency regime (2200-1400 $cm^{-1}$) were separately baseline corrected using a high-pass filter with Gaussian apodization and a cutoff at 30 $cm^{-1}$ for $H_2O$ data and 35 $cm^{-1}$ for $D_2O$ data. A cutoff of 50 $cm^{-1}$ was used exclusively for baseline correction in the high-frequency regime (3500-2900 $cm^{-1}$). The baseline-corrected difference spectra belonging to each dataset were ultimately averaged.

A control measurement was run mixing HU with the employed buffer in the absence of protein. Data were averaged and corrected as described above. The resulting difference spectrum showed only noise in the 2200-1400 $cm^{-1}$ and 3500-2900 $cm^{-1}$ regions, corresponding, respectively, to a maximal peak-to-peak difference of $3x10^{-5}$ and $2x10^{-4}$, or root-mean-square of $0.4x10^{-5}$ and $3x10^{-5}$.

**14. EPR and ENDOR experiments**

EPR and ENDOR spectroscopy(47-49) was used to probe the hydrogen-bonding environment of the DOPA radical. To isolate and characterize the exchangeable proton hyperfine tensors, the ENDOR technique was employed in combination with deuteration of the R2e protein or buffer.

All EPR and ENDOR measurements were performed at the Q-band frequency (approx. 34 GHz) using 500–700 µM solutions of R2e in 1.6 mm quartz tubes. Pulse measurements were carried out at 65 K using a Bruker Elexsys E580 spectrometer equipped with a 150 W TWT amplifier, Bruker EN 5107D2 resonator, Oxford Instruments CF935 continuous-flow helium cryostat and Oxford Instruments MercuryiTC temperature controller. Field-swept EPR spectra were detected via the electron spin echo (ESE) signal. The microwave (MW) $\pi/2$ pulse was 10 ns; the inter-pulse delay $\tau$ was 290 ns. The spectra were pseudo-modulated[49] using either the *fieldmod* function of the EasySpin package[50] or the XEPR software to highlight the spectral features.

Q-band orientation-selective Davies[51] ENDOR spectra were collected with stochastic detection (4) at 65 K using an AR 600 W radiofrequency (RF) amplifier (AR 600A225A). The following MW pulse sequence was used: $\pi_{sel}$−T−π/2−τ−π−τ−echo. The RF pulse was applied during the time interval T ≈ 19 μs and had a length of 17 μs; the selective MW preparation π pulse was 160 ns to resolve $^{1}$H signals with smaller hyperfine couplings; the π/2 and π detection pulses were 10 and 20 ns, respectively; the inter-pulse delay τ was 340 ns.

All EPR and ENDOR simulations were performed using the EasySpin package[50]. When multiple EPR spectra are presented on the same figure, their field axes have been normalized to the same MW frequency.

***EPR experiments***
Figure 3 shows the effect of the protein and buffer deuteration on the EPR line shape, arising from the smaller gyromagnetic ratio of $^{2}$H. A 95% $^{2}$H buffer exchange did not lead to a noticeable narrowing of the DOPA EPR spectrum, as the exchangeable proton hyperfine couplings are generally small as compared to the EPR line width (black and green traces). In contrast, per-deuteration of R2e resulted in a loss of the ~1 mT hyperfine splitting at all three turning points due to the exchange of the beta-proton exhibiting the largest, mostly isotropic hyperfine coupling of $A_{iso}^{\beta 1}$ ≈ 30 MHz (Fig. S13, blue trace)(14, 50). Furthermore, the exchange of protons exhibiting weaker hyperfine interactions lead to a reduction of the EPR line width. Overall, per-deuteration of R2e resulted in the three *g*-components well-resolved already at the Q-band frequency (see marked field positions; in this manuscript we use the naming convention: $g_x > g_y > g_z$).

#### Figures

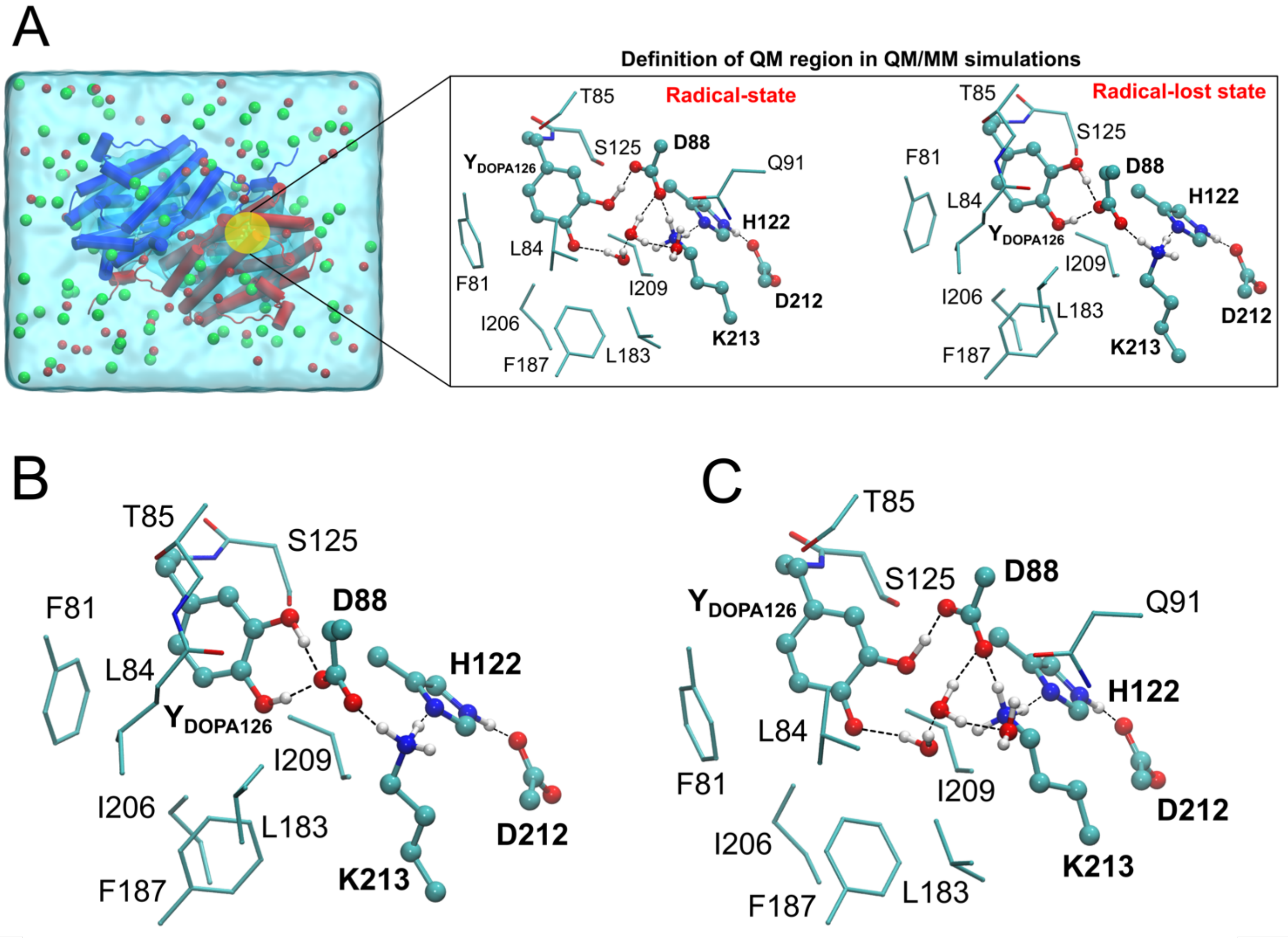

**Fig. S1**. **QM/MM, DFT cluster models, and atomistic MD simulations of R2e. (A)** QM/MM and molecular dynamics (MD) models of the R2e dimer, embedded in a water box with NaCl (green and red spheres). Separate models were created for the radical state (PDB ID: 8bt3) and radical-lost state (PDB ID: 8bt4). *Inset*: The active site, comprising DOPA and the extended hydrogen-bonding network, is highlighted with an orange circle. The QM region used in the QM/MM calculations for simulations of the radical and radical-lost states is shown. **(B-C)** DFT models of R2e active site. The models were constructed from XFEL structure of (**B**) the radical-lost state (PDB ID: 8bt4), and (**C**) the radical state (PDB ID: 8bt3). Key residues involved in the hydrogen-bonding network are highlighted in the CPK representation.

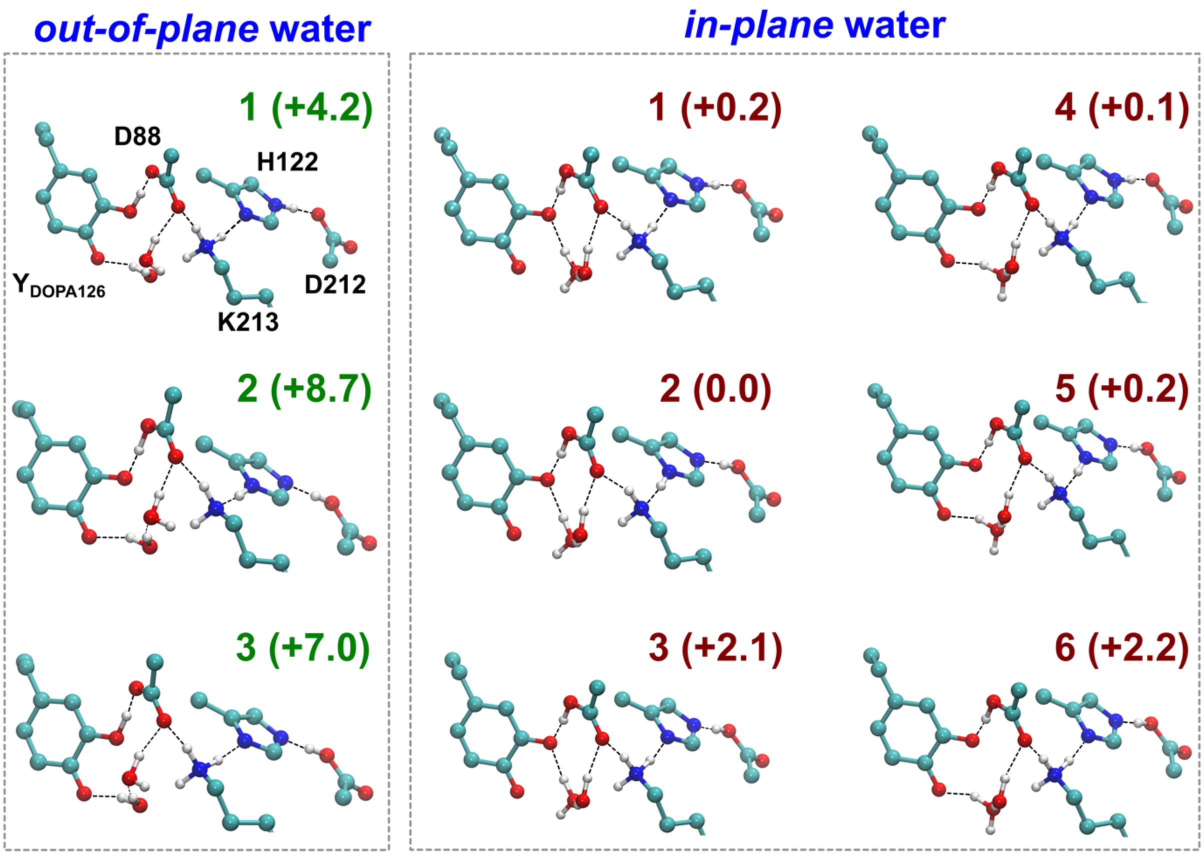

**Fig. S2 DFT models and energetics of the R2e radical state (*S* = 1/2).** Optimized structures with relative electronic energies (in kcal $mol^{-1}$) in the *out-of-plane* and *in-plane* conformations of the water hydrogen bond network. Only the central hydrogen-bond network is shown.

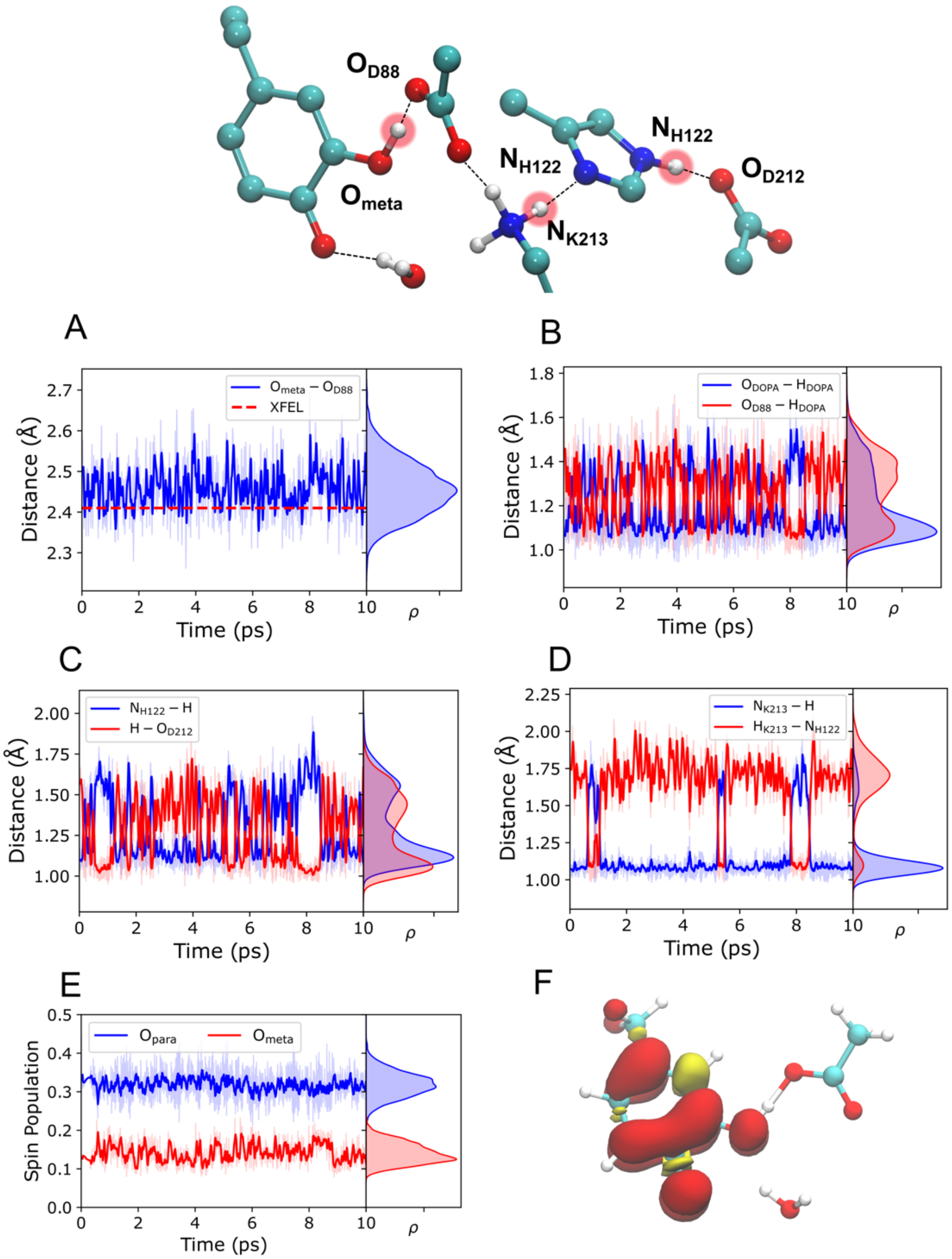

**Fig. S3**. **QM/MM-MD simulations of WT-R2e in the radical state with the *out-of-plane* proximal water molecule.** *Top:* The central hydrogen-bonding network in the radical-lost state, with DOPA forming one hydrogen bond with Asp88. Only central atoms are labelled, while the remaining system is omitted for clarity. **(A)** Dynamics of distances between the meta-oxygen ($O_{meta}$) and para-oxygen ($O_{para}$) of DOPA and Asp88 ($OD_{1/2}$) during the QM/MM-MD simulations. The average $O_{meta}$–$O_{D88}$ from the QM/MM-MD simulations is 2.46 Å, while the horizontal red line corresponds to the distance from the XFEL structure (2.42 Å, PDB ID: 8bt3). Dynamics of the hydrogen-bond distance between **(B)** Asp88 and the meta-OH group of DOPA, **(C)** His122 and Asp212, and **(D)** Lys213 and His122. **(E)** Dynamics of the spin population on the para and meta oxygen atoms of DOPA•, with average spin populations of 0.32 and 0.14, respectively. (**F)** Spin density distribution on DOPA• computed at TPSSh/def2-TZVP-QM/MM level.

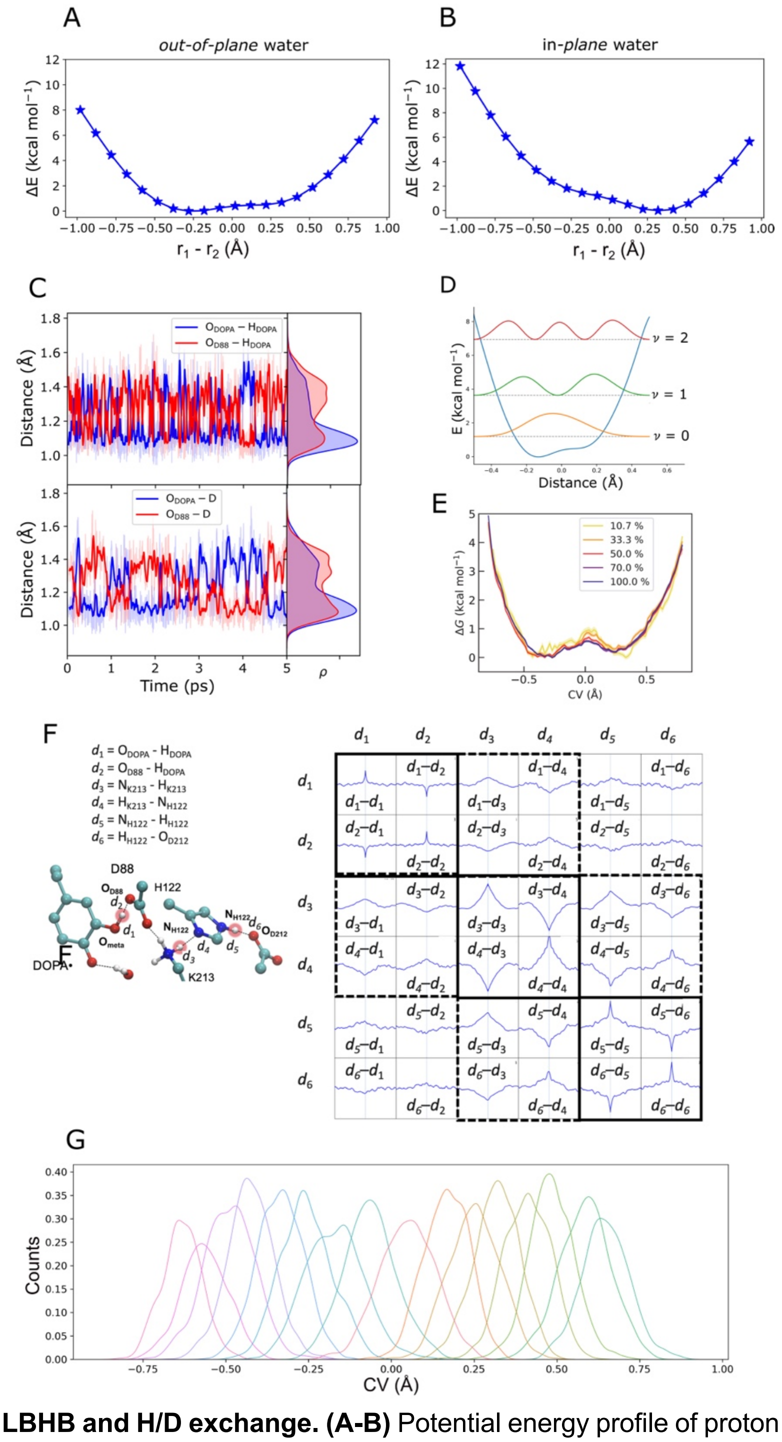

**Fig. S4**. **Energetics of LBHB and H/D exchange. (A-B)** Potential energy profile of proton transfer between $O_{meta}$ of DOPA• and D88 with water in **A,** *out-of-plane* (panel **A**) and **B,** *in-plane* configuration. The potential energy scans were performed at QM/MM (B3LYP/def2-SVP) level. **(C)** Dynamics of hydrogen (top) / deuterium (bottom) bond distance between the $O_{meta}$-Asp88 (Oδ1/Oδ2) during the QM/MM-MD simulations. All hydrogen (deuterium) atoms in this simulation were replaced with deuterium. **(D)** Delocalization of the proton wave function based on numerical solution of the 1D-Schrödinger equation with potential from **A.** (E) Convergence of QM/MM free energy simulations. **(F)** Resonance in the LBHB network form cross-correlation of the $d_1$ – $d_6$ vibrations. A peak in the cross-correlation of the Fourier transformed distance to the frequency domain indicates that the bond distances have resonant vibrations. **(G)** Overlap of umbrella sampling windows for the radical state with the *out-of-plane* water configuration. The collective variable (CV) is defined as $r_1$-$r_2$.

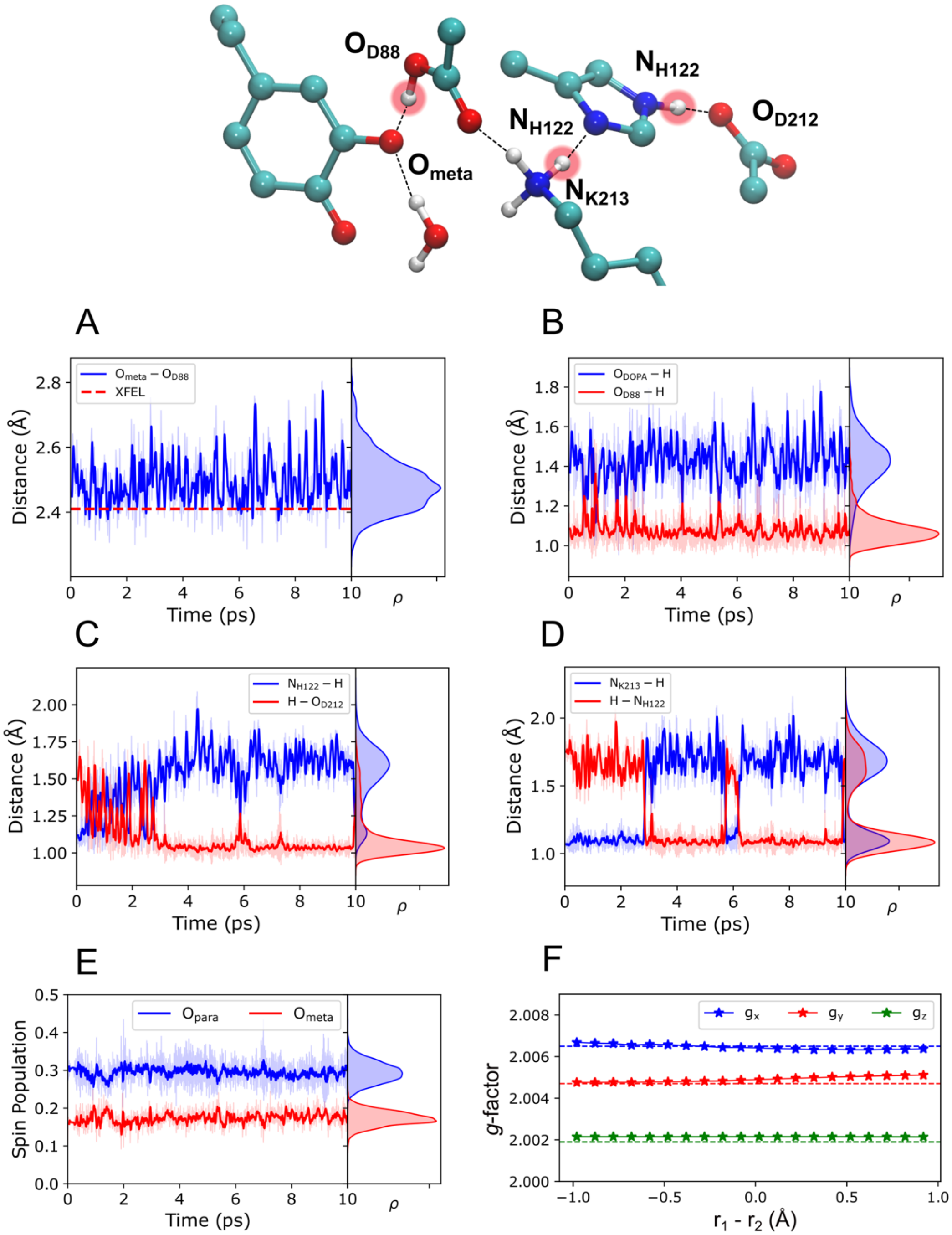

**Fig. S5**. **QM/MM-MD simulations of R2e-WT in the radical state with the *in-plane* proximal water molecule.** *Top:* The central hydrogen-bonding network in the radical-lost state, with DOPA forming one hydrogen bond with Asp88. Only central atoms are labelled, while the remining system is omitted for clarity. **(A)** Dynamics of distances between the meta-oxygen ($O_{meta}$) and para-oxygen ($O_{para}$) of DOPA and Asp88 (OD1/2) during the QM/MM-MD simulations. The average $O_{meta}$–$O_{D88}$ from the QM/MM MD simulations is 2.49 Å, while the horizontal red line corresponds to the distance from the XFEL structure (2.42 Å, PDB ID: 8bt3). Dynamics of the hydrogen-bond distance between **(B)** Asp88 and the meta-OH group of DOPA, **(C)** His122 and Asp212, and **(D)** Lys213 and His122. **(E)** Dynamics of the spin population on the para and meta oxygen atoms of DOPA•, with average spin populations of 0.29 and 0.17, respectively. **(F)** *g*-tensor along the hydrogen-bond distances ($r_1$-$r_2$) for the *in-plane* configuration.

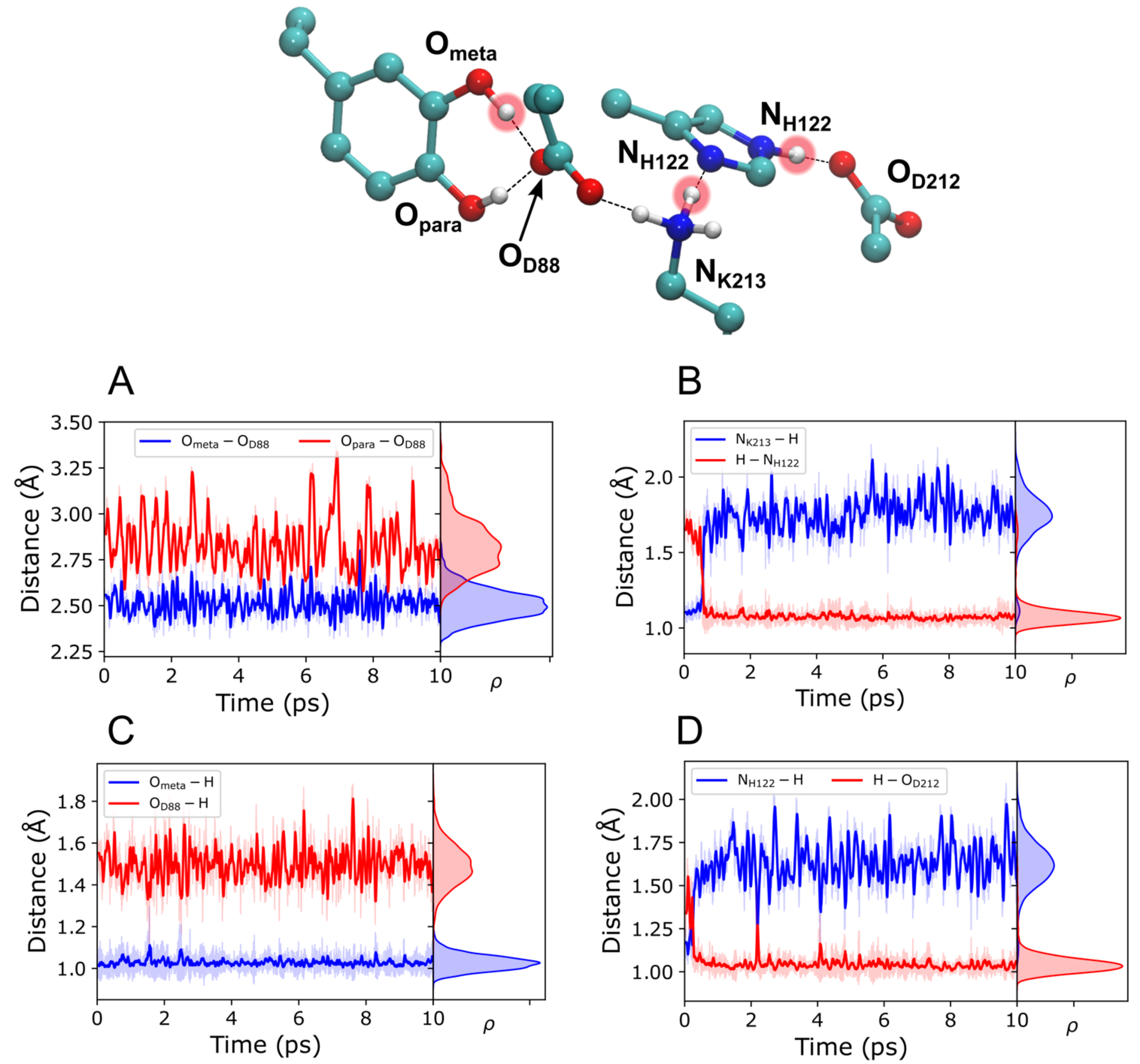

**Fig. S6**. **QM/MM-MD simulations of the radical-lost state with two hydrogen-bonds between DOPA and Asp88 (Conf II).** *Top:* The central hydrogen-bonding network in the radical-lost state, with DOPA forming two hydrogen bonds with Asp88. Only central atoms are labelled, while the remining system is omitted for clarity. **(A)** Dynamics of distances between the meta-oxygen ($O_{meta}$) and para-oxygen ($O_{para}$) of DOPA and Asp88 (OD1/2) during the QM/MM-MD simulations. The average $O_{meta}$–$O_{D88}$ and $O_{para}$–$O_{D88}$ distances are 2.51 Å and 2.83 Å, respectively. Dynamics of the hydrogen-bond distance between **(B)** Lys213 and His122, **(C)** Asp88 and the meta-OH group of DOPA, and **(D)** His122 and Asp212.

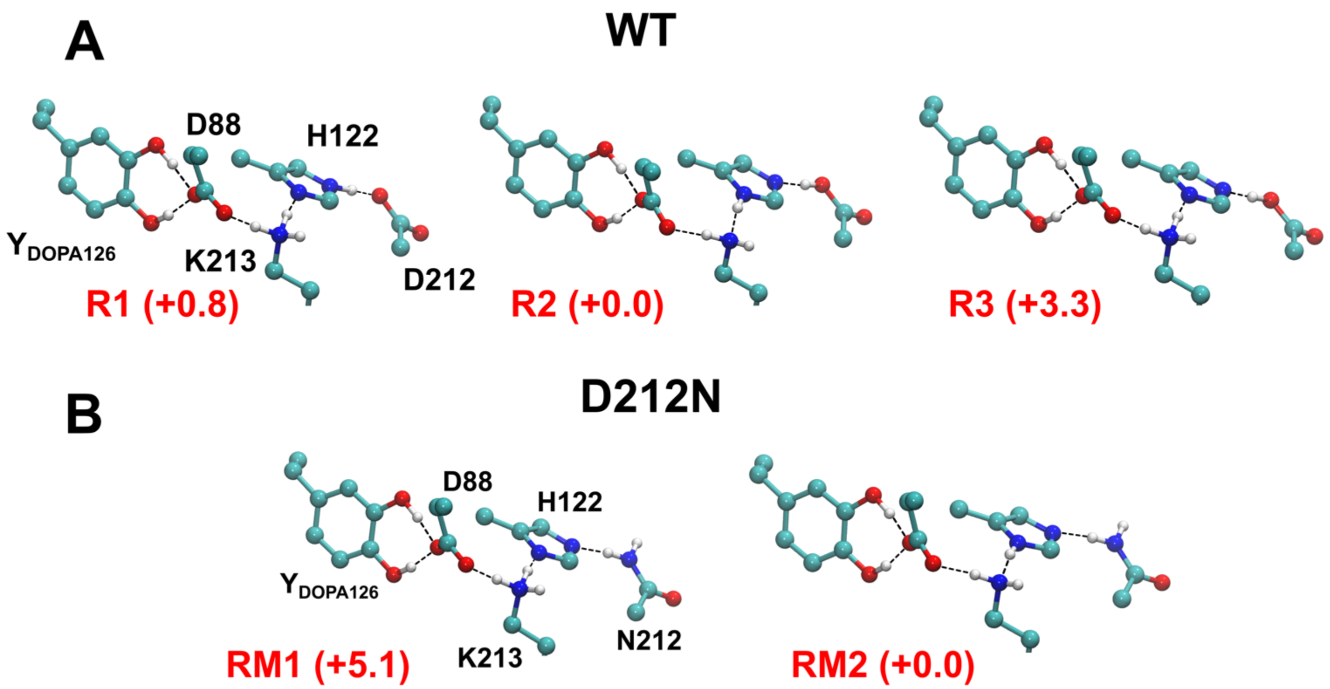

**Fig. S7. DFT models and energetics of the radical-lost state.** Optimized models of **(A)** the WT and **(B)** the D212N variant in the radical-lost state, with relative electronic energies (in kcal $mol^{-1}$). Only the central hydrogen-bonded network is shown. The DFT models of the WT and D212N variant are named R1-R3 and RM1-RM2, respectively.

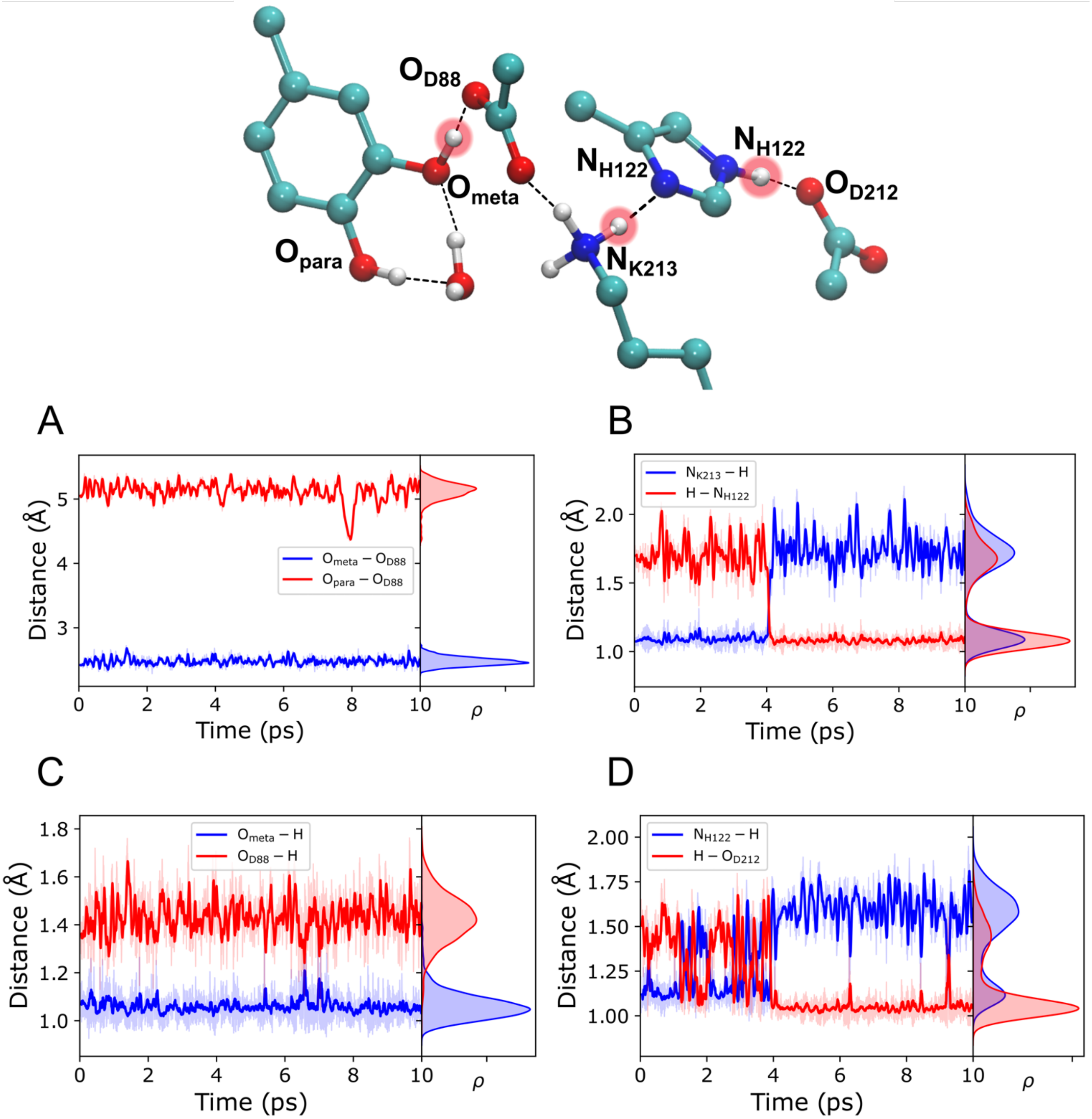

**Fig. S8**. **QM/MM-MD simulations of the radical-lost state with one hydrogen-bonds between DOPA and Asp88 (Conf I).** *Top:* The central hydrogen-bonding network in the radical-lost state, with DOPA forming two hydrogen bonds with Asp88. Only central atoms are labelled, while the remaining system is omitted for clarity. **(A)** Dynamics of distances between the meta-oxygen ($O_{meta}$) and para-oxygen ($O_{para}$) of DOPA and Asp88 (OD1/2) during the QM/MM-MD simulations. The average $O_{meta}$–$O_{D88}$ and $O_{para}$–$O_{D88}$ distances are 2.48 Å and 5.12 Å, respectively. Dynamics of the hydrogen-bond distance between **(B)** Lys213 and His122, **(C)** Asp88 and the meta-OH group of DOPA, and **(D)** His122 and Asp212.

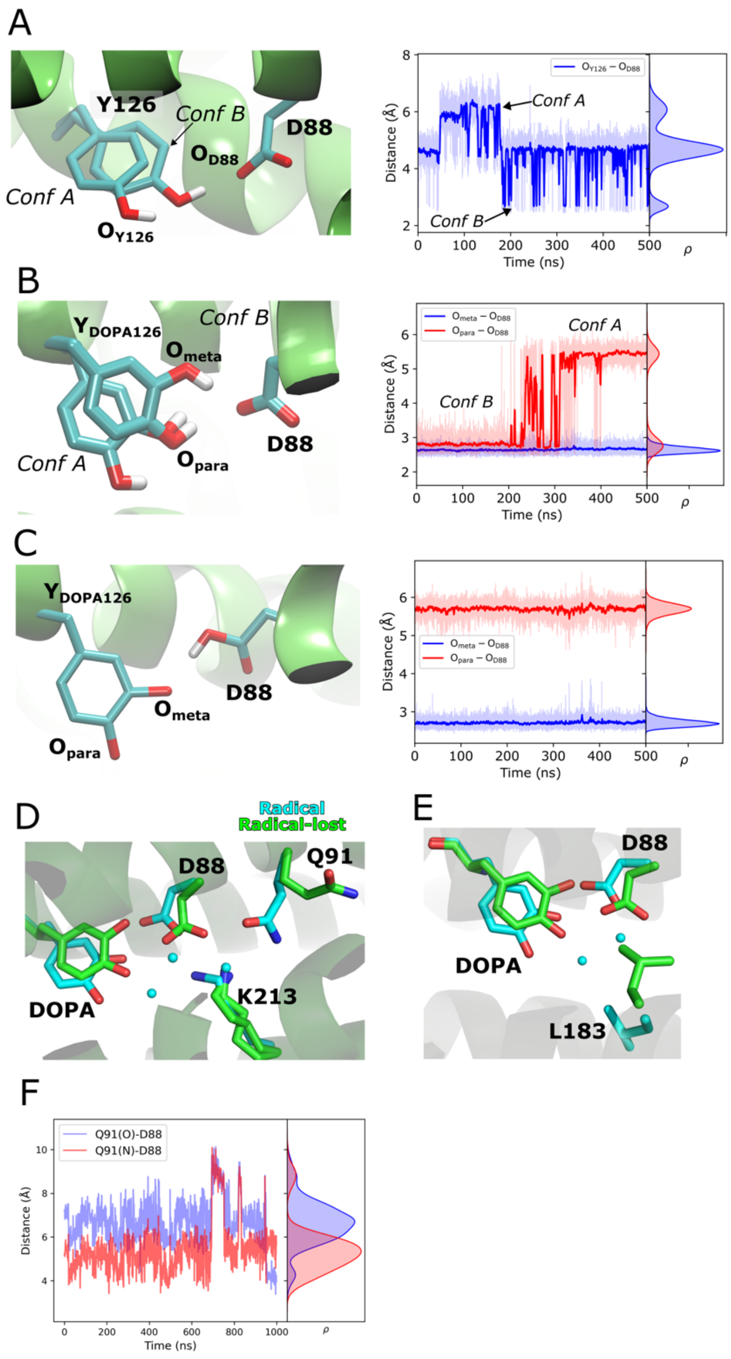

**Fig. S9**. **Classical MD simulations of unactivated Tyr126 and DOPA in the radical and radical-lost states.** **(A)** Conformational dynamics of the Tyr126–Asp88 hydrogen-bond observed in MD simulations (simulation C1, see Table S11). Two possible conformations of Tyr126 are overlaid and correspondingly marked in the distance evolution plot. **(B)** Conformational dynamics of DOPA in the radical-lost state, showing the distance evolution of $O_{meta}$ and $O_{para}$ relative to $O_{D88}$ (simulation C2, see Table S11). Two distinct conformations are highlighted. **(C)** Conformational dynamics of DOPA in the radical state (simulation C3, see Table S11). **(D)** Overlay of XFEL structures of radical (cyan) and radical-lost states (green) highlighting conformational changes in sidechain of Gln91 and DOPA. **(E)** Overlay of XFEL structures of radical (cyan) and radical-lost states (green) highlighting conformational changes in sidechain of Leu183 and DOPA. **(F)** Sidechain conformation of Gln91 in MD simulations of the radical state.

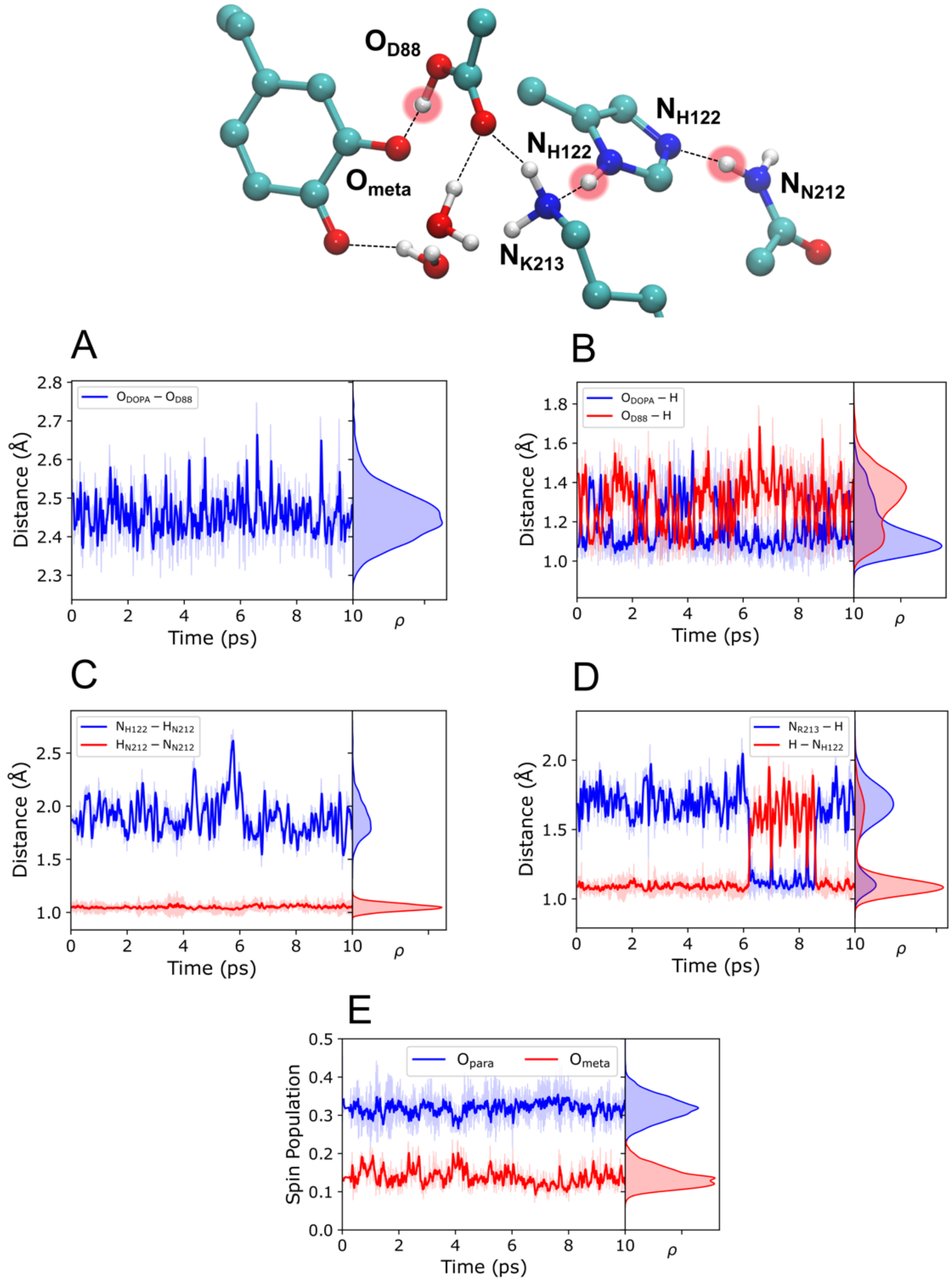

**Fig. S10**. **QM/MM-MD simulations of R2e-D212N in the radical state with the *out-of-plane* proximal water molecule.** *Top:* The central hydrogen-bonding network in the radical-lost state, with DOPA forming one hydrogen bond with Asp88. Only central atoms are labelled, while the remining system is omitted for clarity. **(A)** Dynamics of distances between the meta-oxygen ($O_{meta}$) and para-oxygen ($O_{para}$) of DOPA and Asp88 (OD1/2) during the QM/MM-MD simulations. The average $O_{meta}$–$O_{D88}$ from the QM/MM MD simulations is 2.46 Å. Dynamics of the hydrogen-bond distance between **(B)** Asp88 and the meta-OH group of DOPA, **(C)** His122 and Asp212, and **(D)** Lys213 and His122. **(E)** Dynamics of the spin population on the para and meta oxygen atoms of DOPA•, with average spin populations of 0.32 and 0.14, respectively.

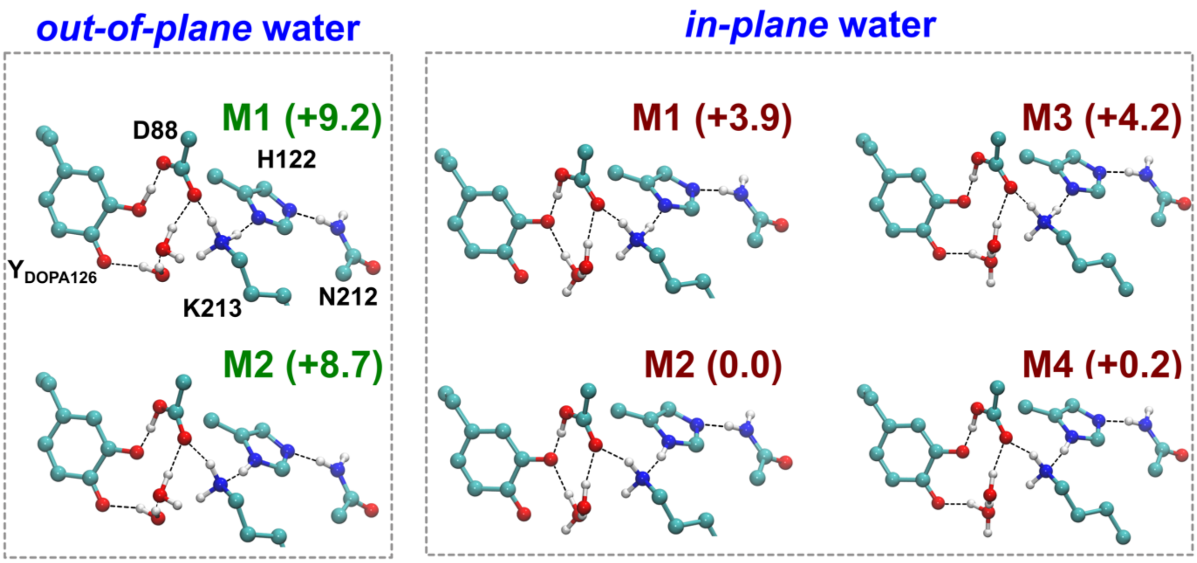

**Fig. S11**. **DFT models and energetics of the D212N variant of R2e in the radical state (S = 1/2).** Optimized structures with relative electronic energies (in kcal $mol^{-1}$) in the *out-of-plane* and *in-plane* conformations of the water hydrogen bond network. Only the central hydrogen-bond network is shown. The DFT models of the D212N variant are named M1-M4.

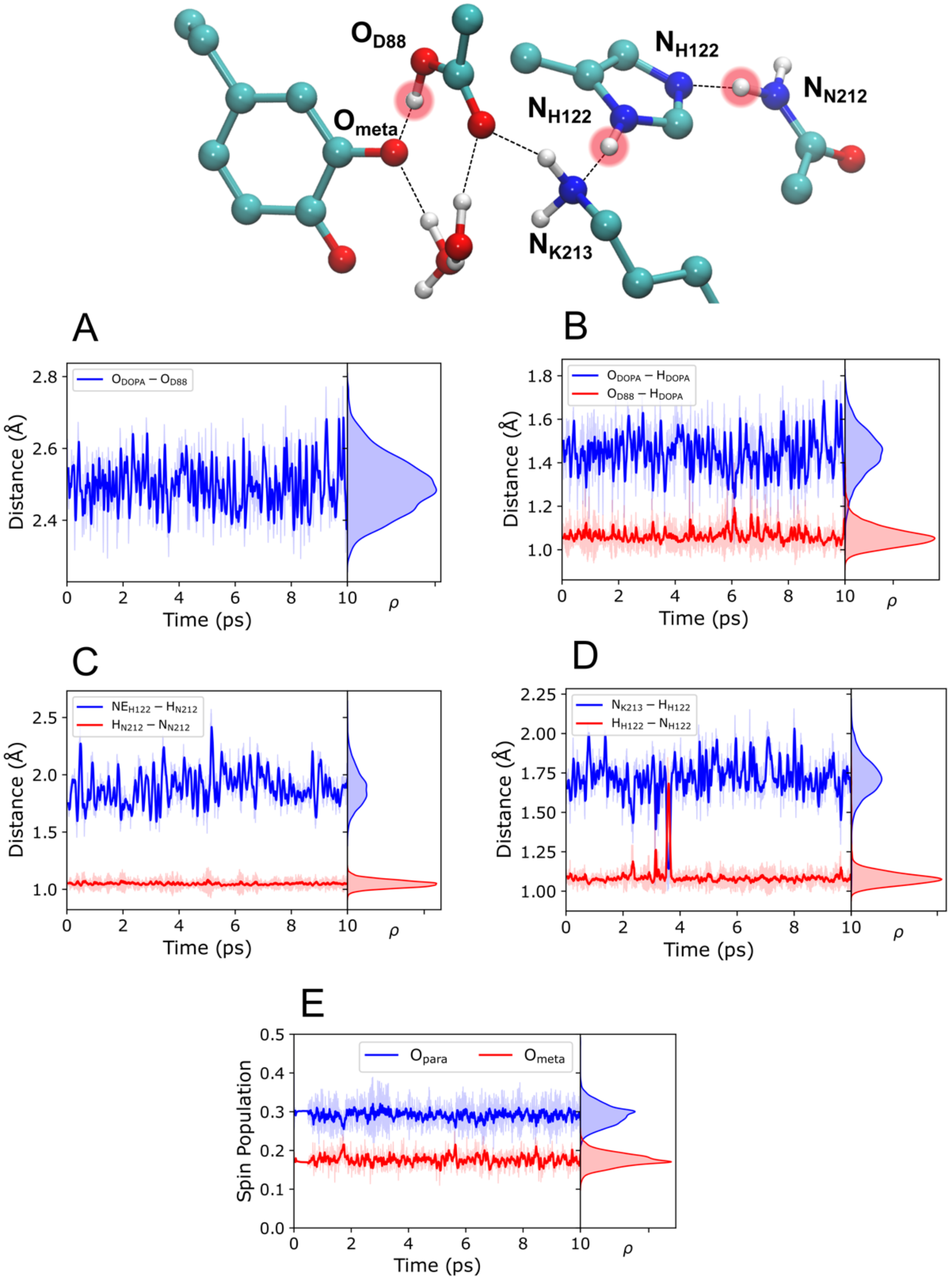

**Fig. S12**. **QM/MM-MD simulations of R2e-D212N in the radical-lost state with the *in plane* proximal water molecule.** *Top:* The central hydrogen-bonding network in the radical-lost state, with DOPA forming one hydrogen bond with Asp88. Only central atoms are labelled, while the remining system is omitted for clarity. **(A)** Dynamics of distances between the meta-oxygen (O$_{meta}$) and para-oxygen (O$_{para}$) of DOPA and Asp88 (OD1/2) during the QM/MM-MD simulations. The average O$_{meta}$–O$_{D88}$ from the QM/MM-MD simulations is 2.50 Å. Dynamics of the hydrogen-bond distance between **(B)** Asp88 and the meta-OH group of DOPA, **(C)** His122 and Asp212, and **(D)** Lys213 and His122. **(E)** Dynamics of the spin population on the para and meta oxygen atoms of DOPA•, with average spin populations of 0.29 and 0.18, respectively.

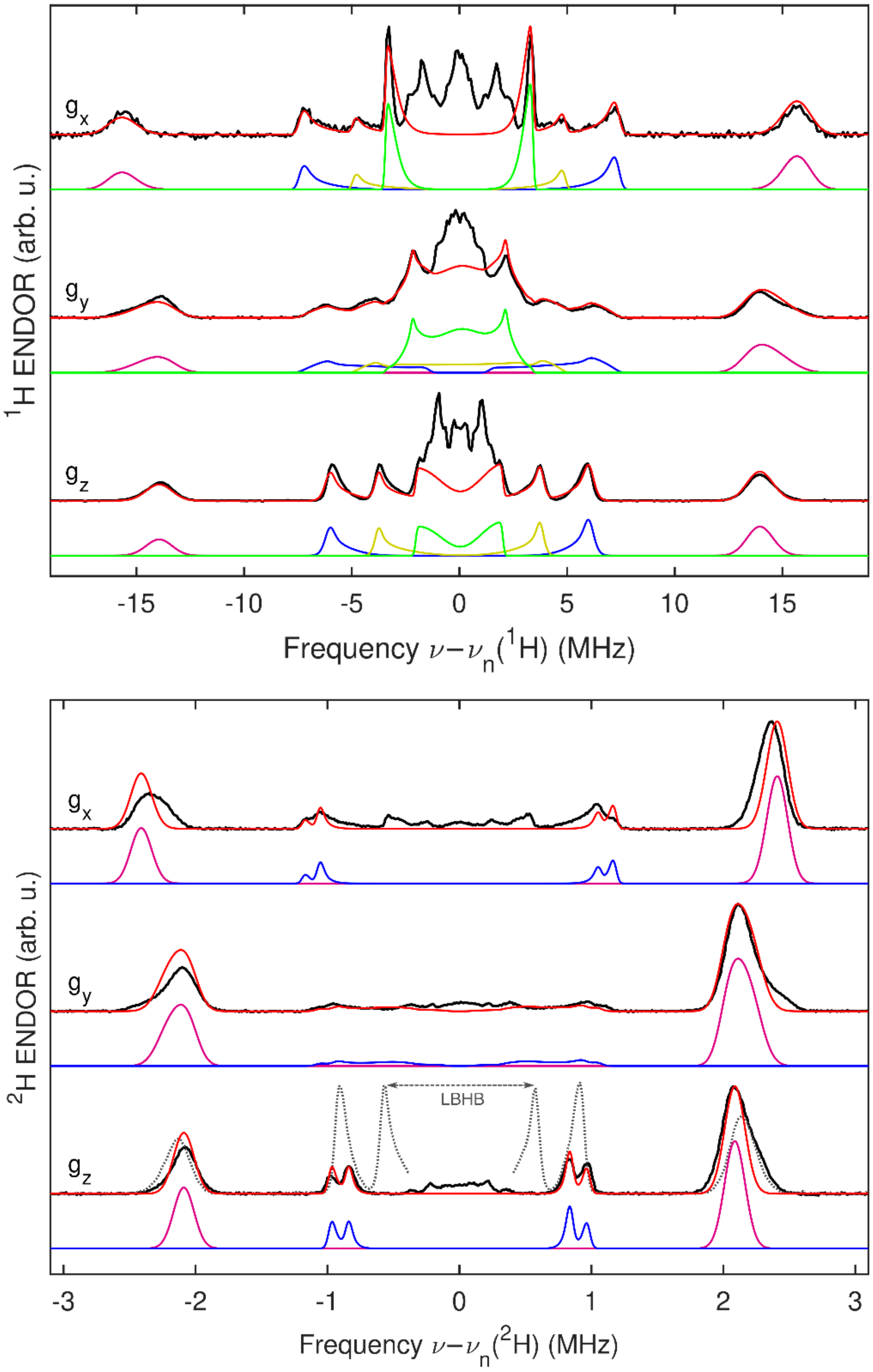

**Fig. S13**. **Orientation-selective Q-band $^{1}H/^{2}H$ ENDOR spectra for the WT R2e radical state.** Top: $^{1}H$ ENDOR of R2e in an $H_2O$ buffer; bottom: $^{2}H$ ENDOR of perdeuterated R2e in an $H_2O$ buffer. The spectra were collected over broad frequency ranges to record all proton/deuteron-related features, including the strongly coupled Cβ-hydron. The dotted line shows an overlaid $^{1}H$ ENDOR trace, scaled to the ratio of gyromagnetic ratios, at the same orientation; the lack of the $^{2}H$ ENDOR feature corresponding to the LBHB deuteron implies a full exchange with buffer protons. Simulations are shown for the most well-resolved hydron signals, with simulation parameters reported in Table S12. A quadruple splitting is clearly resolved for the ring deuteron (bottom, blue trace).

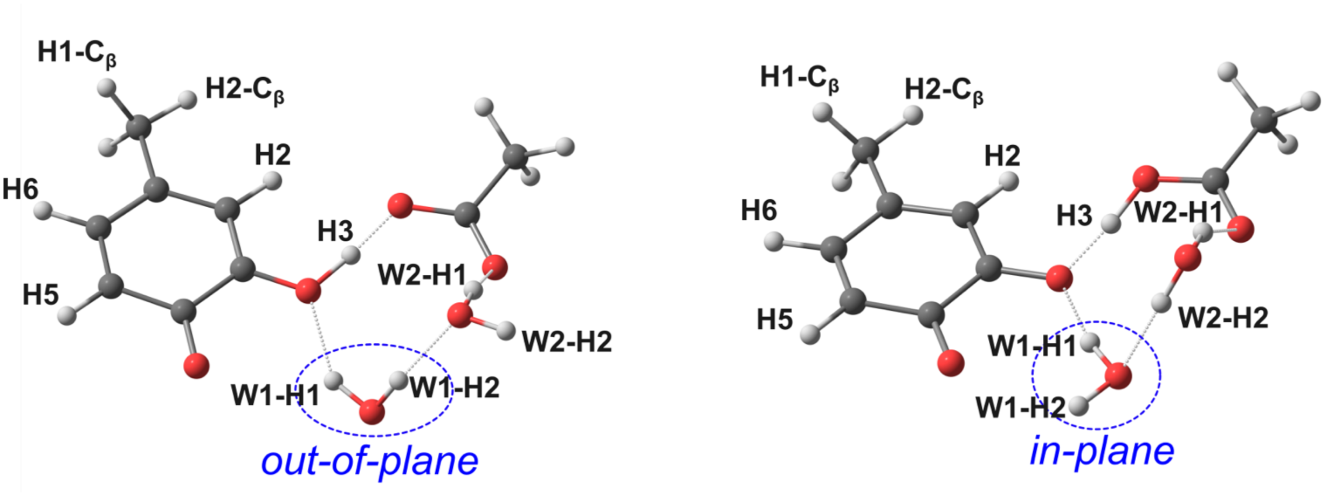

**Fig. S14**. **Nomenclature of the DOPA• ring and $C_\beta$ protons for the hyperfine couplings.** The nomenclature for the DOPA• ring, $C_\beta$, and proximal waters protons is depicted for both *out-of-plane* and *in-plane* water orientation. The small size of the model in the figure is for clarity purposes, actual hyperfine coupling calculations were performed using the large QM region in the QM/MM framework using TPSSh/EPR-II level of theory.

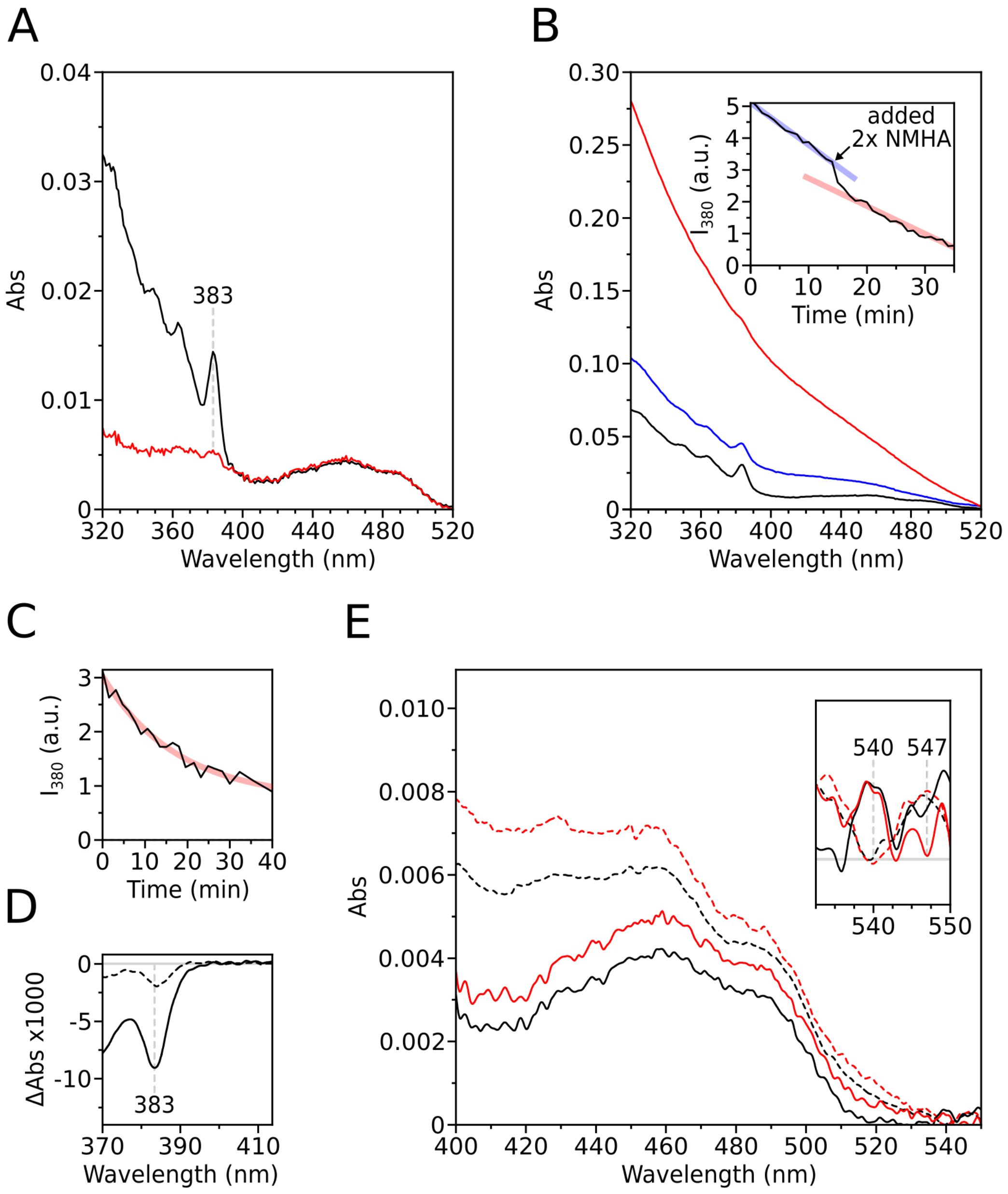

**Fig. S15. Quenching of radical state of WT R2e monitored through UV/Vis spectroscopy**. **(A)** UV/Vis spectra before (black) and after (red) quenching the radical state with 65 mM HU (normalized to OD 1 at 280 nm). **(B)** UV/Vis spectra before (black) and after quenching with 65 mM NMHA (blue) and 115 mM NMHA (red). Spectra normalized to OD 1 at 280 nm. *Inset*: Peak area of the 383 nm peak followed over time and after addition of NMHA (65 mM at t=0, 115 mM total concentration at t=14 min). **(C)** Peak area of the 383 nm peak followed over time after initial addition of 100 mM HU, in $D_2O$ buffer. **(D)** Reaction-induced difference spectra after quenching the radical state with HU of concentration 65 mM in $H_2O$ (solid line) and 100 mM in $D_2O$ (dashed line). **(E)** UV/Vis spectra before (black) and after (red) quenching the radical state with HU, in $H_2O$ (solid lines) and $D_2O$ (dashed lines) buffer. Spectra are offset to yield zero baseline absorbance between 535–545 nm. *Inset*: Magnification of 532.5-550 nm range, showing potential peak shifts upon H/D solvent exchange.

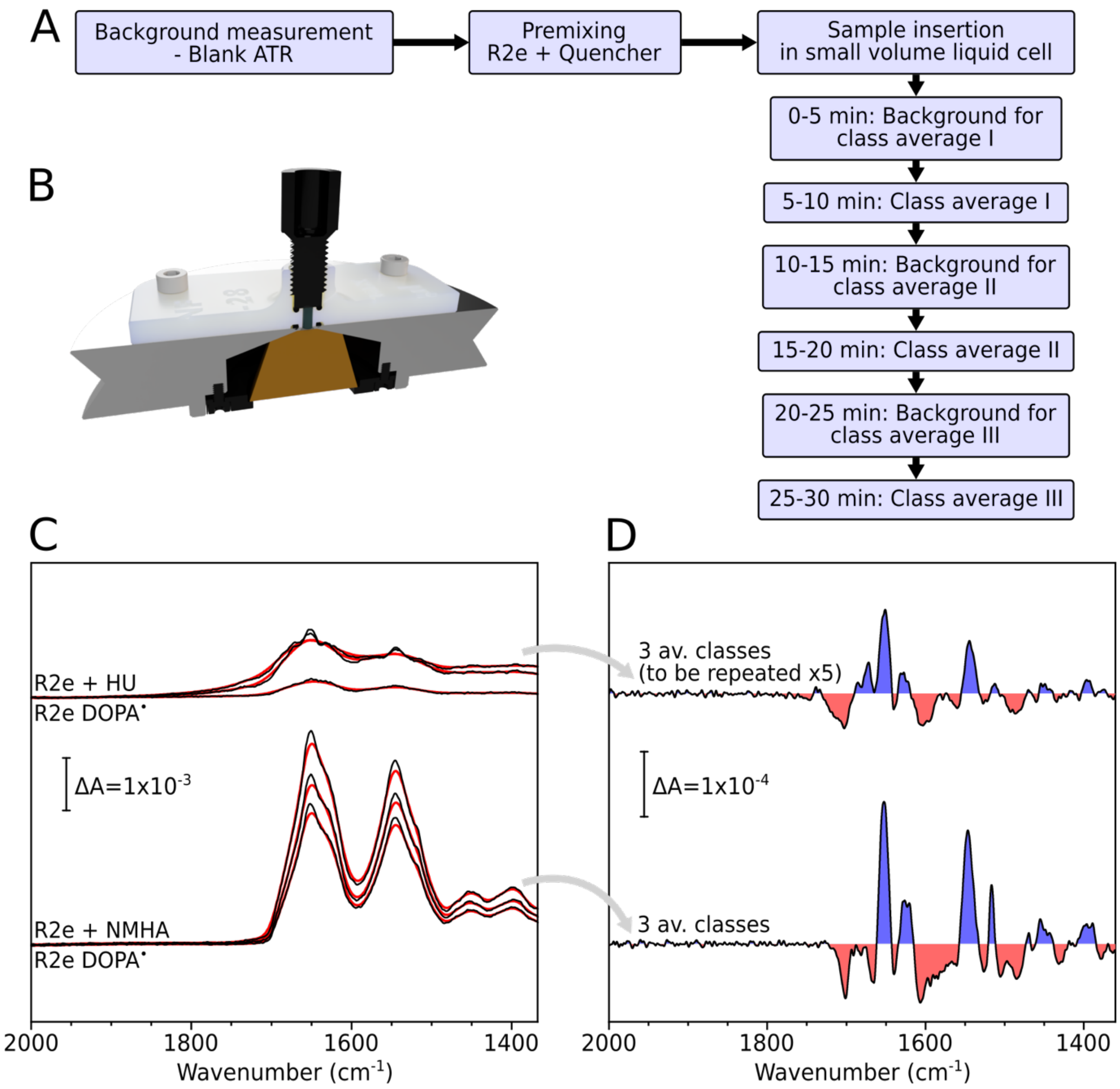

**Fig. S16**. **ATR-FTIR acquisition protocol and baseline correction procedure for reaction-induced difference spectra**. **(A)** Simplified diagram describing the induction of the quenching reaction on the radical state of R2e. **(B)** Rendering of the computer-aided design of the 3D-printed sample holder (small volume liquid cell) mounted onto the Si-ATR cell. Full-section view. **(C)** Difference spectra belonging to each "class" average before baseline correction (black) and averaging; respective baselines are shown as red lines. *Top*: Quenching with HU. *Bottom*: Quenching with NMHA. **(D)** Reaction-induced difference spectra after separate baseline correction and averaging of the three "classes".

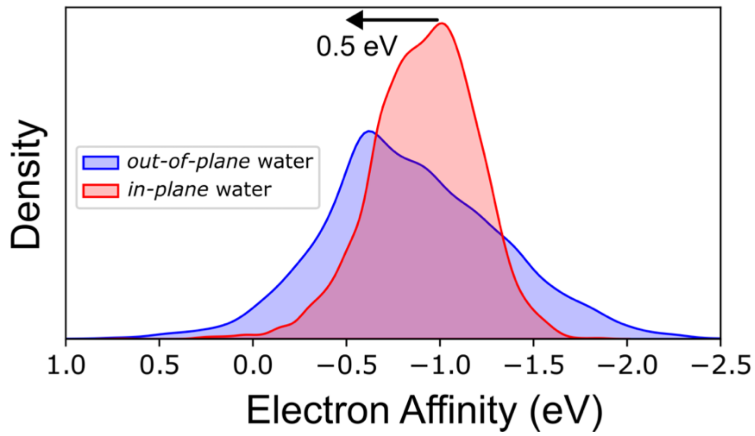

**Fig. S17**. **Redox properties of DOPA•.** Distribution of electron affinity (in eV) of DOPA• in both *out-of-plane* and *in-plane* QM/MM models. Ensembles of vertical electron affinity, respectively, were computed on around 10,000 snapshots extracted from the QM/MM-MD trajectory.

**A**

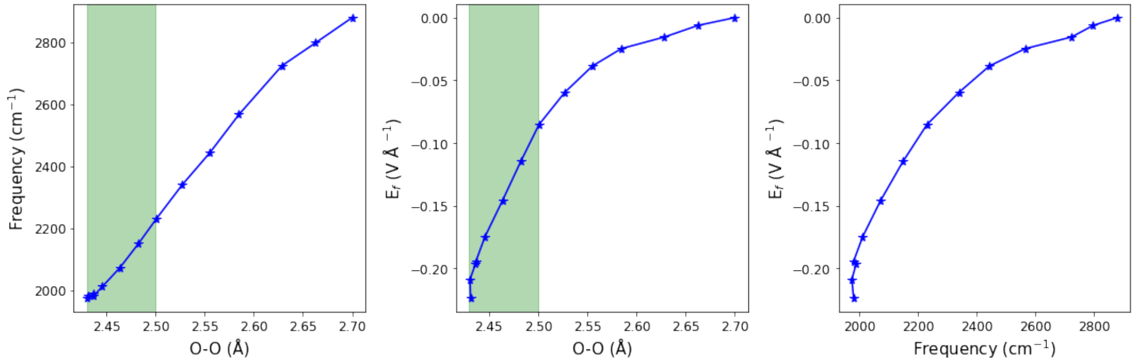

**B**

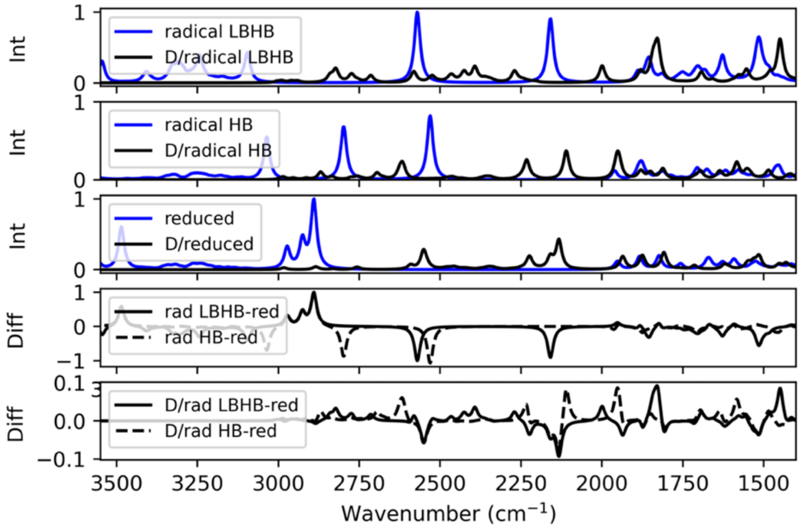

**C**

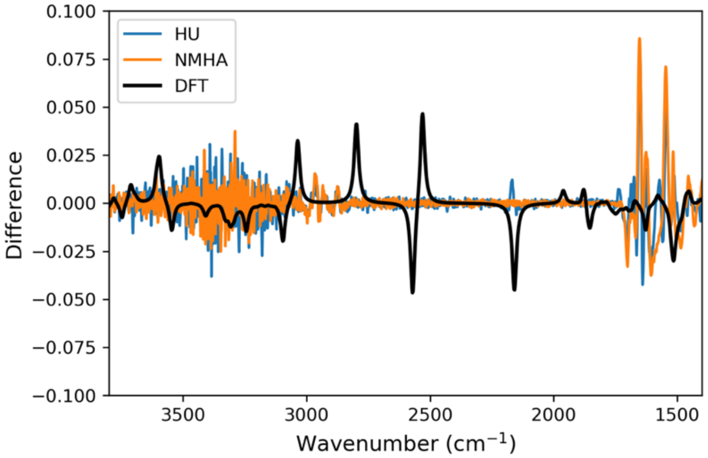

**Fig. S18**. **Electric field and vibrational effects of the LBHB and H/D isotope effects. (A)** Electric field and vibrational effects of the LBHB on DFT models. **(B)** Top: H/D isotope effects. (C) Overlay of experimental and computed difference spectra (see Methods).

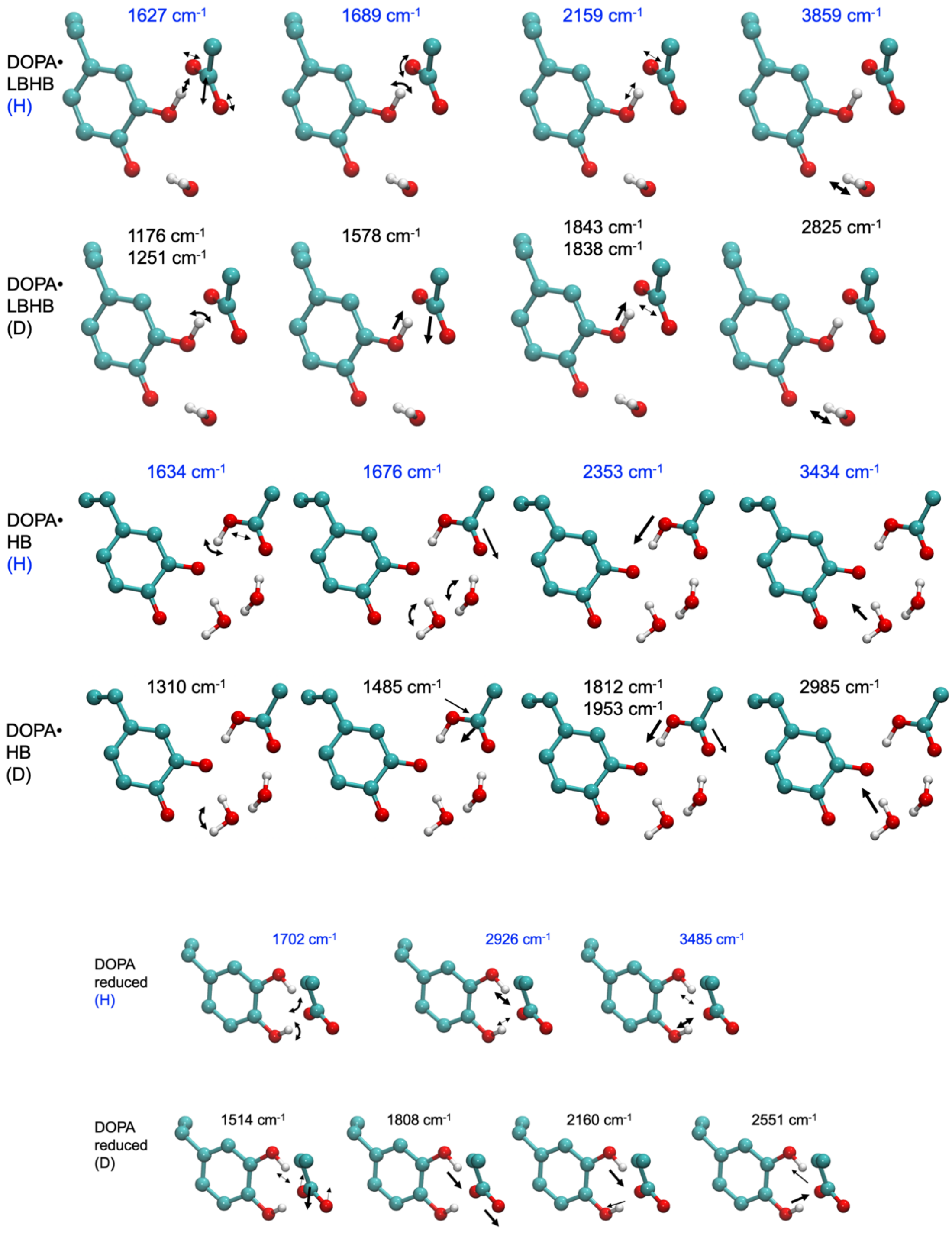

**Fig. S19. Selected vibrational normal modes linked to DOPA/Asp88/proximal water.** The vibrations are empirically shifted to match a characteristic experimental (LBHB-DOPA) vibration at 2159 $cm^{-1}$.

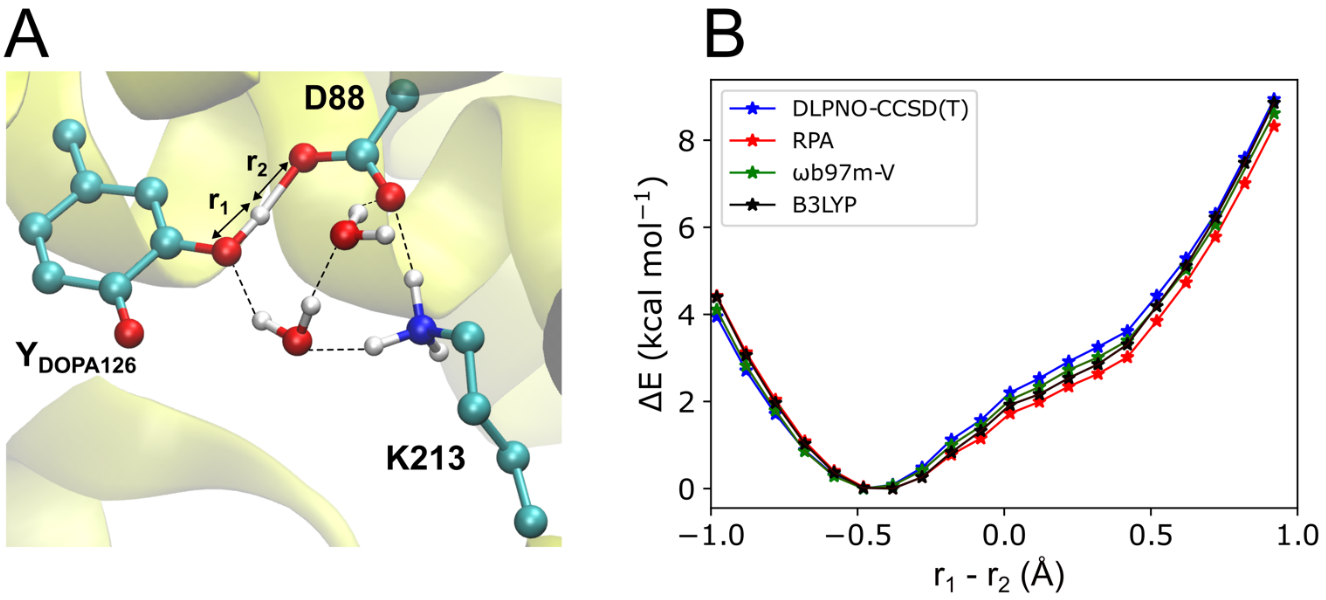

**Fig. S20. Benchmarking the quantum chemical methods for the proton transfer coordinate.** The benchmarking calculations were performed based on QM/MM potential energy scans with a small QM region comprising DOPA•, Asp88, Lys213, and two $H_2O$ molecules, while the remining system was modelled with point charges. **(A)** Reaction coordinate and hydrogen-bonding network. **(B)** Energetics were evaluated using two DFT functionals (B3LYP, ωB97M-V) and two wavefunction-based methods (RPA, DLPNO-CCSD). The calculations performed using def2-TZVPPD basis sets.

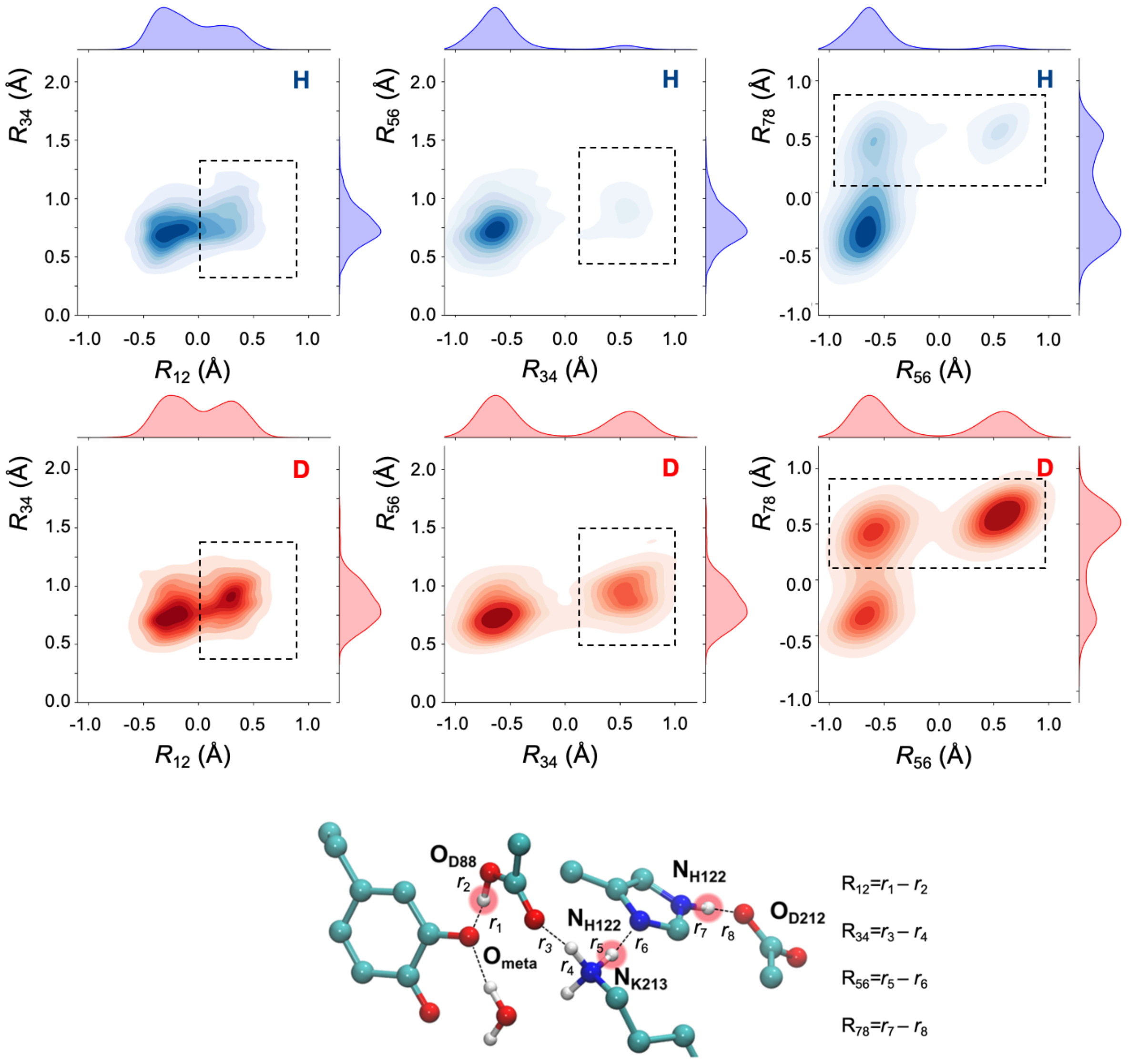

**Fig. S21. H/D isotope effects from QM/MM-MD simulations.** Correlation of hydrogen-bonding distances $R_{12}$-$R_{34}$, $R_{56}$-$R_{34}$, and $R_{56}$-$R_{78}$ from unbiased QM/MM-MD with *Top*: hydrogen and *bottom*: deuterium substituted model. $R_{12}$, measures the difference in proton (deuterium) distance to DOPA ($O_{meta}$) and D88; $R_{34}$ – the difference in D88-K213 proton (D) distances; $R_{56}$ – the difference in $K_{213}$-$H_{122}$ distances; and $R_{78}$ - the difference in $H_{122}$-$O_{212}$ distances, with qualitative differences in the highlighted with dashed squares. The analysis suggests that the proton has a single-well distribution along $R_{12}$ vs. $R_{34}$ relative to the more double-well deuterium profile. The effects extend across the hydrogen-bonded network, as suggested by different populations of, *e.g.*, $R_{56}$ in the protonated and deuterated simulations. The data has been extracted from unbiased QM/MM-MD simulations with a classical description of the proton/deuterium nuclei.

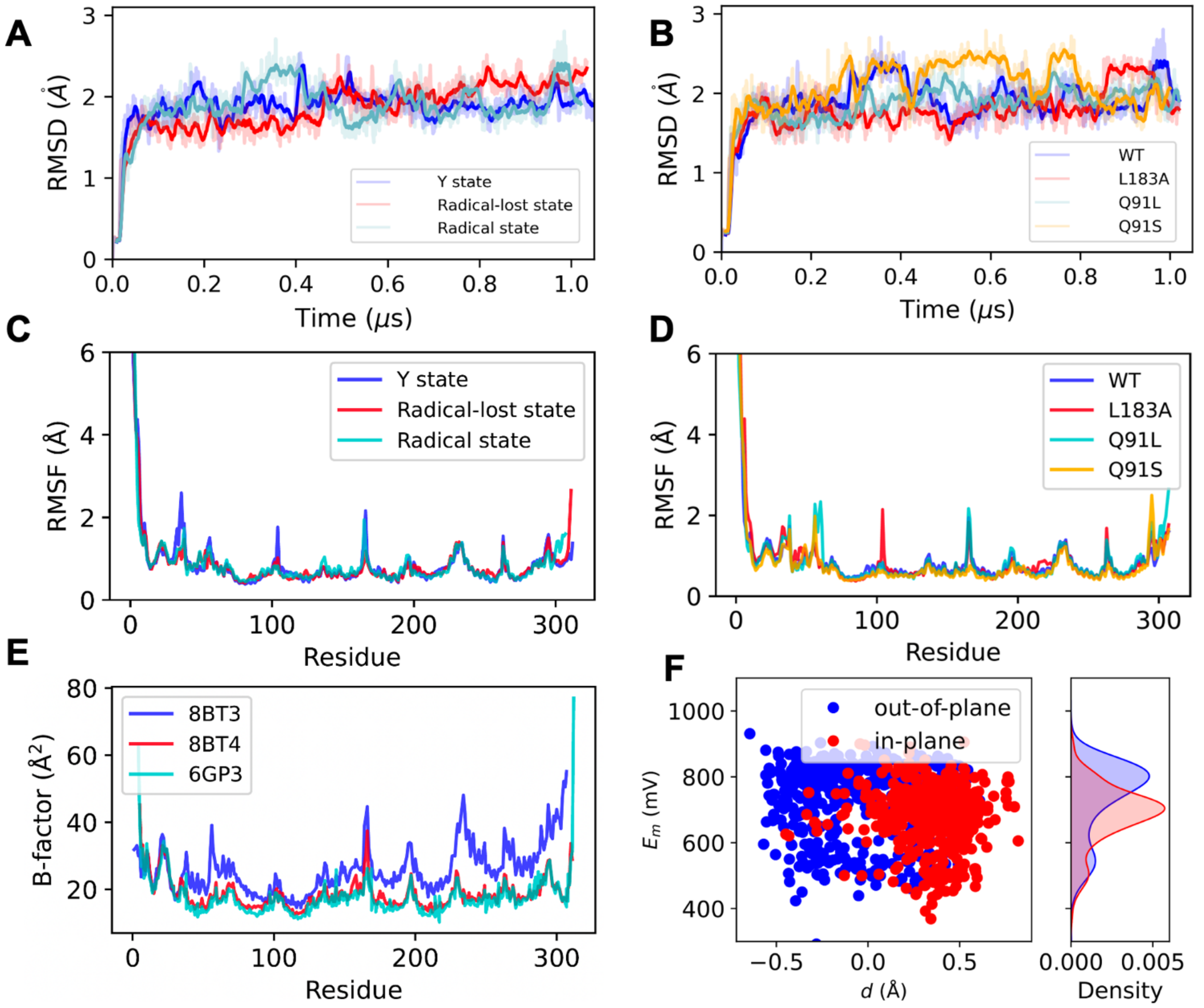

**Fig. S22. Analysis of MD simulations.** A, B, Root-mean-square-deviation (RMSD) of MD simulations of A) WT R2e in different redox states, and B) the L183A, Q91S, and Q91L variants in the radical state. C,D) *Root-mean-square-fluctuation* (RMSF) of C) WT R2e in different redox states, and D) the L183A, Q91S, and Q91L variants in the radical state. E) B-factors from three different x-ray structures (PDB IDs: 8BT3: radical state; 8BT4: radical-lost state; 6GP3: inactive state). F) Correlation of reduction potentials with the water orientation and proton transfer coordinate, $d$=$d_1$-$d_2$ (see Fig. S4E).

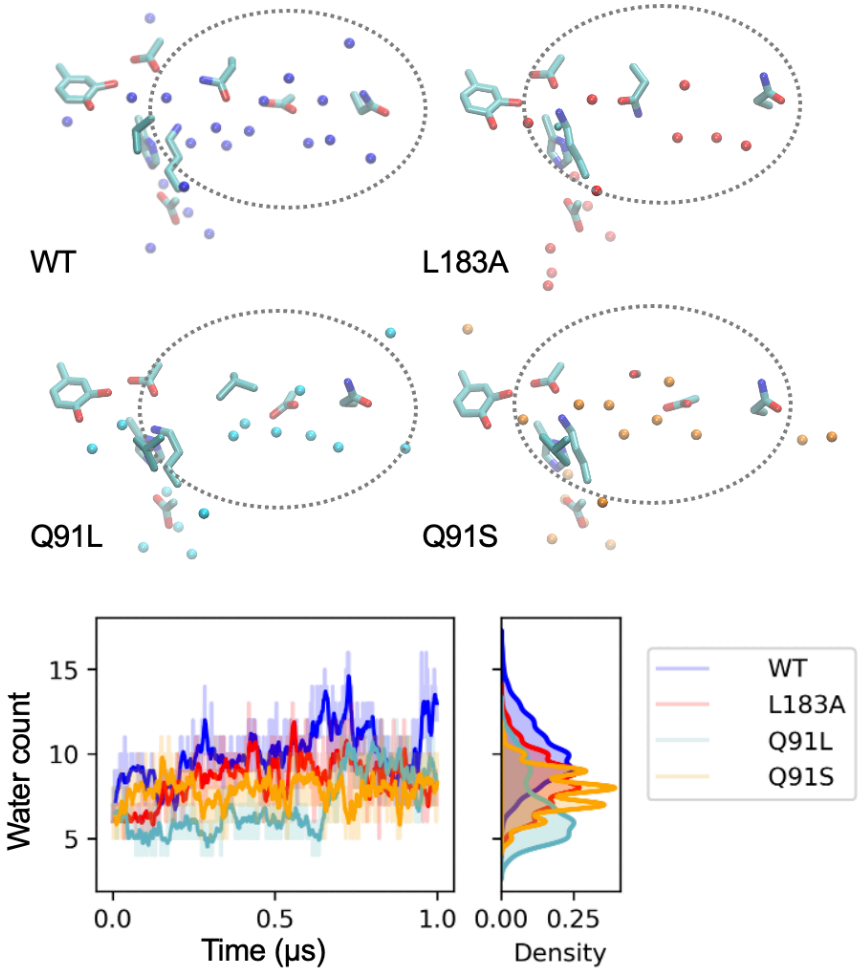

**Fig. S23. MD simulations of the L183A, Q91S, and Q91L variants, and the WT R2e in the radical state.** ***Top***: snapshot from MD simulations, extracted at 1 μs, showing water molecules around the DOPA region. ***Bottom:*** Water count around 4 Å of the active site (residues D88, Q91, D96, Q99, DOPA, K213) during MD simulations.

### Tables

**Table S1. Nuclear quantum effects in R2e.** Impact of nuclear quantum effects on the geometry of the LBHB between DOPA and D88 explored using DFT (classical) and NEO-DFT (quantum) of the proton/deuterium. All distances are reported in units of ångströms.

| **Distance** | **H** (classical) | **H** (quantum) | **D** (quantum) | **Difference** (quantum-classical) |
|---|---|---|---|---|
| O⋯O | 2.56 | 2.52 | 2.52 | -0.04 |
| $O^{DOPA}$-H | 1.02 | 1.06 | 1.05 | +0.04 |
| H-$O^{D88}$ | 1.54 | 1.45 | 1.46 | -0.09 |

**Table S2. Benchmarking of the molecular energetics.** Relative electronic energy (in kcal $mol^{-1}$) of the structural models, with different protonation arrangement along the hydrogen-bonding network (see Fig. S2, *out-of-plane* water models) using different DFT functionals and correlated wavefunction based approaches (RPA and DLPNO-CCSD). DOPA was modeled in the oxidized state. Minimum energy states are marked in bold font.

| Model | TPSS | TPSSh | B3LYP | M06-2X | B2PLYP | ωB97X-D | ωB97M-V | RPA | DLPNO |
|---|---|---|---|---|---|---|---|---|---|
| 1 | **0.00** | **0.00** | **0.00** | 1.62 | 1.58 | 0.22 | 1.25 | 1.86 | 1.22 |
| 2 | 1.11 | 0.73 | 0.24 | **0.00** | **0.00** | **0.00** | **0.00** | **0.00** | **0.00** |
| 3 | 2.17 | 2.15 | 2.49 | 2.15 | 4.36 | 2.61 | 2.67 | 3.01 | 2.38 |

**Table S3.** X-ray data collection and refinement statistics. Statistics for the highest-resolution shell are shown in parentheses.

| | ***Mf* R2e-D212N** |
|---|---|
| **PDB ID** | 9R5L |
| **Data collection statistics** | |
| **Synchrotron/Beamline** | Diamond/i04 |
| **Wavelength [Å]** | 0.95373 |
| **Space group** | C 1 2 1 |
| **Unit cell dimensions a, b, c [Å]** | 141.1, 45.8, 56.8 |
| **Unit cell angles α, β, γ [°]** | 90, 112.2, 90 |
| **Resolution range [Å]** | 43.2-1.7 (1.75 - 1.7) |
| **Unique reflections** | 71037 (5258) |
| **Multiplicity** | 3.6 (3.5) |
| **Completeness (%)** | 97.81 (96.43) |
| **Mean I/sigma (I)** | 9.24 (0.86) |
| **Wilson B-factor [Å$^2$]** | 28.89 |
| **$R_{merge}$ (%)** | 5.42 (121.1) |
| **$R_{meas}$ (%)** | 6.38 (142.6) |
| **$R_{pim}$ (%)** | 3.34 (74.78) |
| **$CC_{1/2}$** | 0.999 (0.596) |
| **CC*** | 1 (0.84) |
| **Refinement statistics** | |
| **Resolution range used in refinement [Å]** | 43.2-1.7 (1.75 - 1.7) |
| **Reflections used in refinement** | 36410 (2728) |
| **Reflections used for $R_{free}$** | 1818 (155) |
| **$R_{work}$ (%)** | 21.11 (45.31) |
| **$R_{free}$ (%)** | 25.67 (48.54) |
| **RMS (bonds)** | 0.006 |
| **RMS (angles)** | 0.75 |
| **Ramachandran favored (%)** | 98.06 |
| **Ramachandran allowed (%)** | 1.94 |
| **Ramachandran outliers (%)** | 0 |
| **Rotamer outliers (%)** | 1.4 |
| **Clashscore** | 3.48 |
| **Protein residues** | 1-309 (of 340) |
| **Average B-factor** | 43.58 |
| **macromolecules** | 43.51 |
| **solvent** | 44.51 |
| **Number of non-H atoms** | 2778 |
| **macromolecules** | 2599 |
| **solvent** | 179 |

**Table S4**. Primers for R2e mutagenesis.

| Mutation | Sequence |
|---|---|
| Q91S for | CATTGTTAGATACAATTAGCGCTACTGTTGGTGATGTG |
| Q91S_rev | CACATCACCAACAGTAGCGCTAATTGTATCTAACAATG |
| Q91L for | CATTGTTAGATACAATTCTGGCTACTGTTGGTGATGTG |
| Q91L rev | CACATCACCAACAGTAGCCAGAATTGTATCTAACAATG |
| L183A for | GCCAGGCTTCTTAGCATATGGAGGCTTCTATTTAC |
| L183A_rev | GTAAATAGAAGCCTCCATATGCTAAGAAGCCTGGC |
| D212N for | ACACTTCAGATATTATTAGATTAATATTAAGAAATAAAGTTATACATAACTACTATAGTGG |
| D212N rev | CCACTATAGTAGTTATGTATAACTTTATTTCTTAATATTAATCTAATAATATCTGAAGTGT |

**Table S5**. **Calculated *g*-tensors for DOPA•.** Comparison of computed and experimental *g*-tensors for optimized DFT cluster model (see *Methods*) with *out-of-plane* and *in-plane* configuration of the proximal water to DOPA•. In addition to this, *g*-tensors of an isolated tyrosyl and DOPA• in gas phase, along with the *g*-tensors of DOPA• from the QM/MM calculations are shown. All computations were performed at TPSSh/EPR-II level of theory. The corresponding *out-of-plane* and *in-plane* water models are shown in Fig. S2.

| **Model** | $g_x$ | $g_y$ | $g_z$ | $g_{iso}$ | $\rho_{O\ (para)}$ | $\rho_{O\ (meta)}$ |
|---|---|---|---|---|---|---|
| **Gas phase** | | | | | | |
| **Tyrosine** | 2.0088924 | 2.0043958 | 2.0021173 | 2.0051352 | 0.39 | - |
| **DOPA** | 2.0064464 | 2.0058673 | 2.0021876 | 2.0048338 | 0.24 | 0.22 |
| | **DFT *out-of-plane* water models** | | | | | |
| **1** | 2.0062955 | 2.0048648 | 2.0020041 | 2.0043882 | 0.29 | 0.13 |
| **2** | 2.0062153 | 2.0051541 | 2.0019949 | 2.0044548 | 0.26 | 0.17 |
| **3** | 2.0062776 | 2.0048619 | 2.0019893 | 2.0043763 | 0.28 | 0.13 |
| **DFT *in-plane* water models** | | | | | | |
| **1** | 2.0063812 | 2.0049907 | 2.0021155 | 2.0044958 | 0.28 | 0.15 |
| **2** | 2.0063598 | 2.0050856 | 2.0021011 | 2.0045155 | 0.27 | 0.16 |
| **3** | 2.0063726 | 2.0050040 | 2.0021194 | 2.0044987 | 0.27 | 0.15 |
| **4** | 2.0057043 | 2.0052051 | 2.0021414 | 2.0043502 | 0.26 | 0.18 |
| **5** | 2.0057968 | 2.0052885 | 2.0021413 | 2.0044088 | 0.25 | 0.18 |
| **6** | 2.0058017 | 2.0050988 | 2.0020117 | 2.0043041 | 0.25 | 0.18 |
| **QM/MM** | | | | | | |
| ***out-of-plane* water** | 2.0066891 | 2.0049078 | 2.0021498 | 2.0045823 | 0.30 | 0.12 |
| ***in-plane* water** | 2.0063389 | 2.0049839 | 2.0021470 | 2.0044899 | 0.28 | 0.15 |
| **Exp.** | 2.0065 | 2.0047 | 2.0019 | 2.0044 | - | - |

**Table S6. DFT computed hyperfine coupling constants for the *out-of-plane* water orientation.** The hyperfine coupling constants (in MHz) for the DOPA• ring, Cβ, and proximal water protons (*out-of-plane* model) are tabulated along with the corresponding Euler angles (in degrees). The calculations were performed using a QM/MM approach at the TPSSh/EPR-II level of theory. The nomenclature of the protons is shown in Fig. S14.

| | $A_x$ | $A_y$ | $A_z$ | $A_{iso}$ | α (degrees) | β (degrees) | γ (degrees) |
|---|---|---|---|---|---|---|---|
| **H1-Cβ** | 26.82 | 27.54 | 31.92 | 28.759 | 85.4 | 21.3 | -128.5 |
| **H2-Cβ** | -0.87 | -1.17 | 2.84 | 0.267 | -64.1 | 20.5 | 82.3 |
| **H2** | 1.11 | 3.05 | 7.31 | 3.826 | 81.9 | 12.4 | -80.9 |
| **H3** | -9.20 | 7.36 | -12.12 | -4.652 | 94.6 | 32.9 | -92.2 |
| **H5** | -14.01 | -3.74 | -16.82 | -11.523 | 83.1 | 23.8 | -85.1 |
| **H6** | -4.47 | 1.97 | -4.13 | -2.208 | -125.0 | 9.0 | 123.8 |
| **W1-H1** | -2.78 | 3.59 | -0.39 | 0.142 | -93.8 | 44.2 | 89.8 |
| **W1-H2** | -0.92 | 1.87 | -0.60 | 0.117 | -79.6 | 38.8 | 89.5 |
| **W2-H1** | -0.57 | 1.09 | -0.52 | -0.001 | -35.5 | 27.2 | 53.7 |
| **W2-H2** | -0.30 | 0.60 | -0.27 | 0.011 | -55.5 | 30.5 | 75.6 |

**Table S7. DFT computed hyperfine coupling constants for the *in-plane* water orientation.** The hyperfine coupling constants (in MHz) for the DOPA• ring, Cβ, and proximal water protons (*in-plane* model) are tabulated along with the corresponding Euler angles (in degrees). The calculations were performed using a QM/MM approach at the TPSSh/EPR-II level of theory. The nomenclature of the protons is shown in Fig. S14.

| | $A_x$ | $A_y$ | $A_z$ | $A_{iso}$ | α (degrees) | β (degrees) | γ (degrees) |
|---|---|---|---|---|---|---|---|
| **H1-Cβ** | 19.04 | 19.63 | 23.78 | 20.819 | -103.3 | 18.0 | 138.3 |
| **H2-Cβ** | -0.61 | -0.81 | 2.76 | 0.445 | 49.0 | 23.5 | -89.7 |
| **H2** | -1.39 | 0.62 | 4.56 | 1.265 | 91.8 | 8.4 | -93.1 |
| **H3** | -5.14 | 7.59 | -5.91 | -1.150 | -108.9 | 30.1 | 105.1 |
| **H5** | -11.57 | -2.29 | -13.28 | -9.047 | -82.6 | 17.6 | 84.1 |
| **H6** | -7.07 | 0.39 | -8.55 | -5.076 | 82.2 | 16.6 | -81.2 |
| **W1-H1** | -3.92 | -0.72 | 4.44 | -0.068 | -92.2 | 25.6 | 93.8 |
| **W1-H2** | -1.99 | -1.27 | 3.26 | 0.0001 | -91.8 | 37.5 | 92.7 |
| **W2-H1** | -0.66 | 1.25 | -0.59 | -0.0005 | 36.5 | 38.8 | -59.8 |
| **W2-H2** | -0.87 | 1.56 | -0.66 | 0.0085 | 62.5 | 44.6 | -84.0 |

**Table S8. Variation of the DOPA• *g*-tensor and $^{1}H$ HFC along the LBHB proton transfer coordinate.** Computed *g*-tensors of DOPA• as a function of the low-barrier hydrogen bond (LBHB) proton transfer coordinate (*R*). In addition, the hyperfine coupling constants (MHz) and the corresponding Euler angles (degrees) are shown for the LBHB proton. All structures used in the calculations were obtained from the QM/MM-based potential energy scan. All EPR calculations were performed at the TPSSh/EPR-II level of theory.

| $R=r_1-r_2$ | $g_x$, $g_y$, $g_z$ | $A_x$, $A_y$, $A_z$, $A_{iso}$ | α, β, γ |
|---|---|---|---|
| 0.92 | 2.0065055, 2.0052967, 2.0021514 | -2.65, 4.69, -2.51, -0.1528 | -90.3, 25.6, 80.7 |
| 0.82 | 2.0064928, 2.0052708, 2.0021521 | -2.93, 5.13, -2.86, -0.2191 | -62.5, 30.0, 50.3 |
| 0.72 | 2.0065088, 2.0052464, 2.0021527 | -3.25, 5.62, -3.33, -0.3186 | -133.5,38.0, 132.5 |
| 0.62 | 2.0065124, 2.0052179, 2.0021536 | -3.64, 6.05, -3.85, -0.4765 | -115.8, 30.7, 112.3 |
| 0.52 | 2.0065203, 2.0051896, 2.0021540 | -4.11, 6.52, -4.49, -0.6939 | -109.0, 29.8,105.2 |
| 0.42 | 2.0065368, 2.0051578, 2.0021530 | -4.67, 6.94, -5.26, -0.9963 | -106.2, 31.3, 102.5 |
| 0.32 | 2.0065605, 2.0051192, 2.0021539 | -5.29, 7.21, -6.13, -1.4021 | -102.9, 30.9, 99.4 |
| 0.22 | 2.0065801, 2.0050823, 2.0021535 | -5.97, 7.46, -7.09, -1.8696 | -101.5, 31.6, 98.2 |
| 0.12 | 2.0066075, 2.0050386, 2.0021537 | -6.67, 7.58, -8.11, -2.3984 | -99.8, 32.7, 96.6 |
| 0.02 | 2.0066370, 2.0050030, 2.0021532 | -7.40, 7.64, -9.20, -2.9878 | -99.0, 32.3, 96.0 |
| -0.08 | 2.0066666, 2.0049606, 2.0021532 | -8.03, 7.65, -10.18, -3.5179 | -98.1, 33.1, 95.2 |
| -0.18 | 2.0067186, 2.0049365, 2.0021515 | -8.60, 7.52, -11.10, -4.0603 | -97.7, 33.4, 95.0 |
| -0.28 | 2.0067659, 2.0049142, 2.0021501 | -9.07, 7.32, -11.92, -4.5586 | 96.4, 33.2, -94.0 |
| -0.38 | 2.0068137, 2.0048920, 2.0021497 | -9.32, 7.25, -12.45, -4.8409 | 96.5, 33.2, -94.1 |
| -0.48 | 2.0068303, 2.0048704, 2.0021492 | -9.49, 7.12, -12.80, -5.0552 | 96.5, 33.3, -94.1 |
| -0.58 | 2.0069042, 2.0048633, 2.0021488 | -9.54, 7.02, -13.02, -5.1814 | 96.4, 33.4, -94.1 |
| -0.68 | 2.0069503, 2.0048550, 2.0021486 | -9.56, 6.88, -13.14, -5.2732 | 95.6, 33.4, -93.4 |
| -0.78 | 2.0069779, 2.0048438, 2.0021487 | -9.47, 6.74, -13.06, -5.2643 | 96.3, 33.5, -94.2 |
| -0.88 | 2.0070193, 2.0048448, 2.0021494 | -9.42, 6.62, -13.00, -5.2686 | 95.2, 32.9, -93.1 |
| -0.98 | 2.0070333, 2.0048374, 2.0021492 | -9.36, 6.54, -12.98, -5.2644 | 95.8, 33.0, -93.9 |

**Table S9. Modeled QM region in the QM/MM simulations.** Residues modeled in the QM region of the QM/MM simulations.

| QM/MM model | QM region | Number of QM atoms |
|---|---|---|
| Q1 | Phe81, Leu84, Thr85, Asp88, His122, Ser125-$Y_{DOPA}$126, Leu183, Phe187, Ile206, Ile209, Asp212 and Lys213. | 161 |
| Q2 | Phe81, Leu84, Thr85, Asp88, Gln91, His122, Ser125-$Y_{DOPA}$126, Leu183, Phe187, Ile206, Ile209, Asp212, Lys213 and 3 $H_2O$ | 182 |
| Q3-Q5 | Phe81, Leu84, Thr85, Asp88, Gln91, His122, Ser125-$Y_{DOPA}$126, Leu183, Phe187, Ile206, Ile209, Asp212, Lys213 and 3 $H_2O$ | 181 |
| Q6-Q7 | Phe81, Leu84, Thr85, Asp88, Gln91, His122, Ser125-$Y_{DOPA}$126, Leu183, Phe187, Ile206, Ile209, Asn212, Lys213 and 3 $H_2O$ | 182 |
| Q8 | Trp52, His122, Asp212, Lys213 | 55 |

**Table S10**. **List of QM/MM MD simulations of various redox state of R2e.** Simulations were performed for the WT and D212N variant, with the corresponding protonation network marked around DOPA. DOPA and DOPA• refer to the "reduced" and "oxidized" states, respectively, and with the *in-plane*/*out-of-plane* water orientation indicated.

| QM/MM-MD | PDB | Type | Initial protonation configuration | Simulation length |
|---|---|---|---|---|
| | | **Radical-lost state** | | |
| Q1 | 8bt4 | WT | DOPA-$Asp_{88}$-$Lys_{213}$-$His_{122}$(ε)-$Asp_{212}$ | 10 ps |
| Q2 | 8bt3 | WT | DOPA-$Asp_{88}$-$Lys_{213}$-$His_{122}$(ε)-$Asp_{212}$ | 10 ps |
| | | **Radical state** | | |
| Q3 | 8bt3 | WT | DOPA•-$Asp_{88}$-$Lys_{213}$-$His_{122}$(ε)-$Asp_{212}$ ***in-plane*** **water** | 10 ps |
| Q4 | 8bt3 | WT | DOPA•-$Asp_{88}$-$Lys_{213}$-$His_{122}$(ε)-$Asp_{212}$ ***in-plane*** **water (deuterated)** | 5 ps |
| Q5 | 8bt3 | WT | DOPA•-$Asp_{88}$-$Lys_{213}$-$His_{122}$(ε)-$Asp_{212}$ ***out-of-plane*** **water** | 10 ps |
| Q6 | 8bt3 | D212N | DOPA•-$Asp_{88}$*-$Lys_{213}$*-$H_{122}$(δ)-$Gln_{212}$* ***in-plane*** **water** | 10 ps |
| Q7 | 8bt3 | D212N | DOPA•-$Asp_{88}$-$Lys_{213}$-$His_{122}$(δ)-$Gln_{212}$* ***out-of-plane*** **water** | 10 ps |
| **Total** | | | | **65 ps** |

**Table S11. List of classical MD simulations of various redox states of R2e.** Simulations were performed for the WT, with the corresponding protonation network marked around DOPA. DOPA and DOPA• refer to the "radical-lost" ("reduced") and "radical" states, respectively. Non-standard protonation state of the residues along the network are marked with asterisk (*). The identity of the ε and δ tautomer of $H_{122}$ is shown in the parenthesis.

| MD | PDB | Type | Protonation Network | Simulation length |
|---|---|---|---|---|
| | | **Unactivated state** | | |
| C1 | 6gp3 | WT | $Y_{126}$-$Asp_{88}$-$Lys_{213}$-$His_{122}$(ε)-$Asp_{212}$ | 2 x 500 ns |
| | | **Radical-lost state** | | |
| C2 | 8bt4 | WT | DOPA-$Asp_{88}$-$Lys_{213}$-$His_{122}$(ε)-$Asp_{212}$ | 2 x 500 ns |
| | | **Radical state** | | |
| C3 | 8bt3 | WT | DOPA•-$Asp_{88}$-$Lys_{213}$-$His_{122}$(ε)-$Asp_{212}$ | 2 x 500 ns |
| | | **Radical state** | | |
| C4 | 8bt3 | Q91S | DOPA•-$Asp_{88}$-$Lys_{213}$-$His_{122}$(ε)-$Asp_{212}$ | **2 x 500 ns** |
| | | **Radical state** | | |
| C5 | 8bt3 | Q91L | DOPA•-$Asp_{88}$-$Lys_{213}$-$His_{122}$(ε)-$Asp_{212}$ | **2 x 500 ns** |
| | | **Radical state** | | |
| C6 | 8bt3 | L183A | DOPA•-$Asp_{88}$-$Lys_{213}$-$His_{122}$(ε)-$Asp_{212}$ | **2 x 500 ns** |
| **Total** | | | | **2 x 3.0 μs** |

**Table S12. Experimentally fitted hyperfine parameters for simulation of the ENDOR spectra.** The hyperfine tensor values are in MHz (experimental error within 0.1 MHz). To minimize the number of fitting parameters, all hyperfine tensors were assumed collinear with the *g*-tensor.

| | $A_X$ | $A_Y$ | $A_Z$ |
|---|---|---|---|
| **H1-Cβ** | 32.0 | 26.5 | 27.8 |
| **D1-Cβ**[b] | 31.7 | 26.3 | 27.1 |
| **H5** | −14.8 | $\lvert A_Y \rvert < 3$[a] | −12.1 |
| **D5**[b] | −14.7 | —[a] | −11.8 |
| **H3** | −9.84 | 6.00 | −7.52 |
| **D3**[b] | −10.53 | —[a] | −8.12 |
| **H6** | 6.9 | 4.3 | −4.1 |

[a] The $A_Y$ values of H5/D5 and D3 could not be determined due to a high number of broadened overlapping ENDOR signals in this field region (in Fig. S13 $A_Y = -2.8$ MHz was used). [b] Hyperfine values of deuterons (D1-Cβ, D3, D5) were multiplied by 6.514 (ratio of the gyromagnetic ratios) to simplify the comparison to the proton values.

**Table S13. List of protonation states used in MD simulations.** p$K_a$ values of relevant titratable residues assessed with H++ server (pypka (51, 52) with standard settings), as well as by performing with the PBE/MC approach, at ε=10 for the explicit protein environment and ε=80 for the implicit water environment using APBS/Karlsberg+ (36). The rest of the titratable residues were modelled in their reference protonation state.

| **Residue** | **PDB ID:8BT3 monomer** | **PDB ID:8BT4 dimer** | **PDB ID:6GP3 dimer** | **H++ server (pypKa)** |
|---|---|---|---|---|
| Asp-88 | 2.3 | 3.0 | | 2.6 |
| His-122 | <2 ε tautomer | <2 ε tautomer | <2 ε tautomer | n. a. |
| His-143 | 7.2 | 9.3 | 10.5 | 6.9 |
| Asp-212 | 1.5 | <2 | <2 | 0.8 |
| Lys-213 | 21 | >20 | >20 | n. a. |
| His-216 | 6.8 ε tautomer | 10.8 | 10.6 | 6.7 |

#### Legends for movies

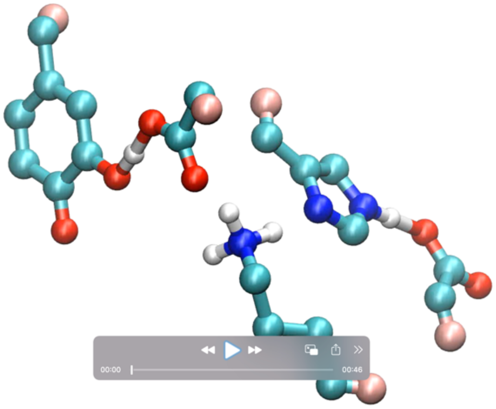

**Movie S1**. Proton transfer dynamics in the LBHB network around DOPA• from QM/MM-MD simulations.

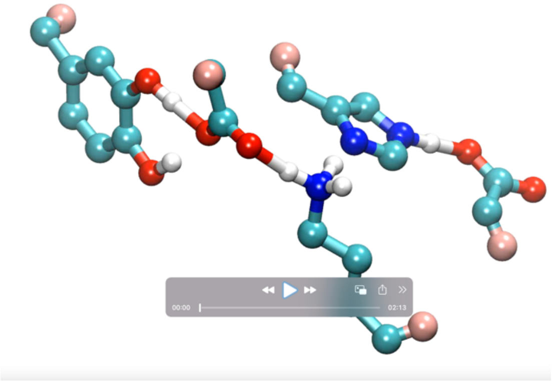

**Movie 2**. Proton transfer dynamics in the hydrogen-bonding network dynamics in radical-lost state from QM/MM-MD simulations.
